# Microbe-derived indole tunes organ-function and links microbe metabolites to biological ageing

**DOI:** 10.1101/2023.01.24.525337

**Authors:** Peter Yuli Xing, Anusha Jayaraman, George Wei Zhang, Katherine Ann Martin, Llanto Elma Faylon, Staffan Kjelleberg, Scott A. Rice, Yulan Wang, Adesola T. Bello, Elaine Holmes, Jeremy K Nicholson, Luke Whiley, Sven Pettersson

## Abstract

To investigate the underlying molecular mechanisms on how the gut microbe metabolite, indoles, regulate host organ growth and function, germ-free male mice were mono-colonized with indole-producing wildtype *Escherichia coli* or tryptophanase-encoding *tnaA* knockout mutant indole-non-producing *E. coli*. The indole mutant *E. coli* recipient mice exhibited significant multiorgan decline and growth retardation combined with catabolism and energy deficiency despite increased food intake compared to control mice. In addition, indole mutant mice displayed malfunctional intestine, enlarged caecum, reduced numbers of colonic enterochromaffin cells and reduced circulating serotonin levels, resulting in reduced gut motility, diminished digestion, and lower energy harvest. Furthermore, indole mutant mice also displayed decreased expression of *Kcnj12* gene, suggesting reduced excitability of enteric neurons thus adding to intestinal dysfunctional phenotype. In conclusion, indoles are necessary to maintain adult metabolic homeostasis across multiple organs in vivo. Impairment of indole levels results in multiorgan functional decline suggesting a mechanism whereby gut microbe metabolites may regulate biological ageing and thus increase the risk for disease.

**Graphical summary:** 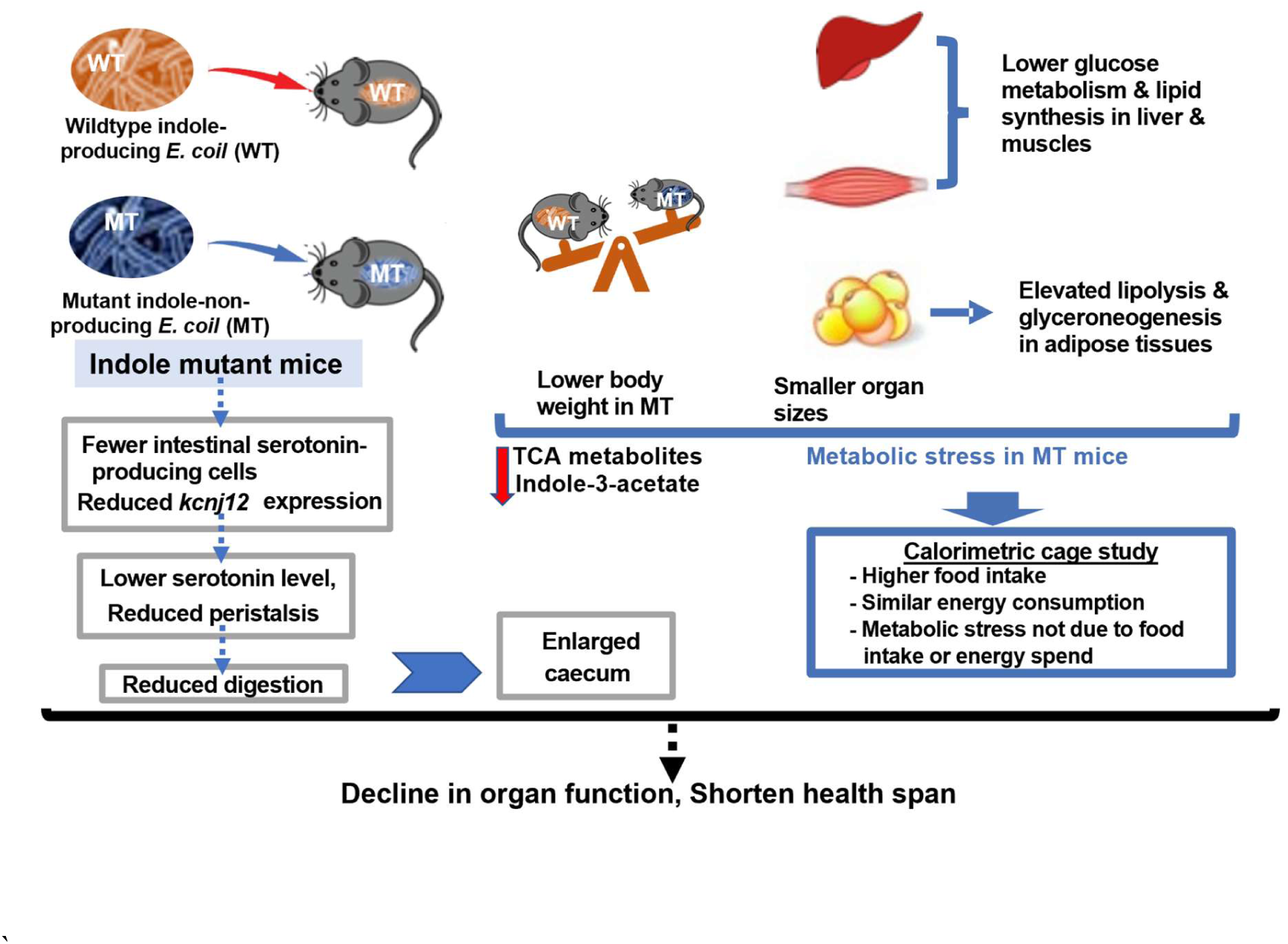

## INTRODUCTION

Host-gut microbiome interactions occur via bidirectional communication through diverse molecules, including host-derived antimicrobial peptides and microbe-generated short-chain fatty acids (SCFAs) (Rooks & Garrett, 2016). Indole(s) are tryptophan metabolites derived from gut microbe metabolism through tryptophanase (encoded by *tnaA*), exemplified by *Bacteroides thethaiotamicron* and *Escherichia coli* (*E. coli*) (Abdul Rahim et al, 2019). Germ-free (GF) mice have very low levels of indoles due to a lack of gut microbiota (Wikoff et al, 2009). In contrast, indole concentrations in healthy rodents and humans have been reported to be about 80 nmol/g and 2.6 mM, respectively (Darkoh et al, 2015; Dong et al, 2020). Indoles have been shown to extend the health span of flies, worms, and rodents based on their enhanced ability to maintain higher composite health scores, reproductive ability, and more youthful genetic expression profiles (Sonowal et al, 2017). Indole and indole-3-acetate (I3A) are important regulators of metabolic homeostasis; indole regulates glucose metabolism through colonic enteroendocrine L cells *in vitro* (Chimeral et al., 2014), whereas I3A inhibits hepatic gluconeogenesis and attenuates cytokine-mediated lipogenesis (Smith et al., 1979). Interestingly, the systemic indole concentration inversely correlates with body mass index and obesity in human subjects and mice (Virtue et al, 2019; Ma et al, 2020). These observations are consistent with a model whereby gut microbe-indole production may impact organ homeostasis relevant to the ageing body.

Reduced indole levels in adult humans with metabolic syndrome/obesity and rats fed with high-fat diet suggest that a correlation exists between gut microbial indole production and metabolic homeostasis (Jennis et al, 2018; Liu et al, 2021). We took a reductionistic approach and introduced a mutation in *E. coli* by removing the tryptophanase-encoding gene *tnaA* (Δ*tnaA*), thereby eliminating the conversion of tryptophan into indoles by these bacteria. GF mice were colonized with wildtype (indole-producing) or Δ*tnaA* (indole-non-producing) *E. coli* (hereafter referred to as WT mice and MT mice, respectively). This approach left the eukaryotic genetic phenotype intact, enabling us to monitor the interplay between organ function and gut microbes with and without indoles.

## RESULTS

### Absence of gut microbial indole production reduces host organ weight of several metabolic organs

MT mice had a constant reduction in body weight throughout the 13-week monitoring period compared to WT mice (**Figure 1A and 1B**, **Supplemental Figure 1A**). Detailed analyses showed significant differences between WT and MT mice in the gross weights of the main metabolic organs: liver, interscapular brown adipose tissue (BAT), epididymal white adipose tissue (WAT), and hindlimb fast glycolytic muscles gastrocnemius (GS) and quadriceps (QC) (**Figure 1C** to **G**). The weights of two other hindlimb muscles, tibialis anterior (TA, fast twitch) and soleus (slow twitch), also tended to be reduced but not significantly (**Figure 1B, C**). Surprisingly, no signs of atrophy were observed in the tissues tested (**Supplemental Fig. 3 and 4A**). Notably, these multiple changes in different organs were observed in mice with an intact genetic configuration thus better reflecting the situation of an accelerated organ decline or accelerated ageing.

**Figure 1.**
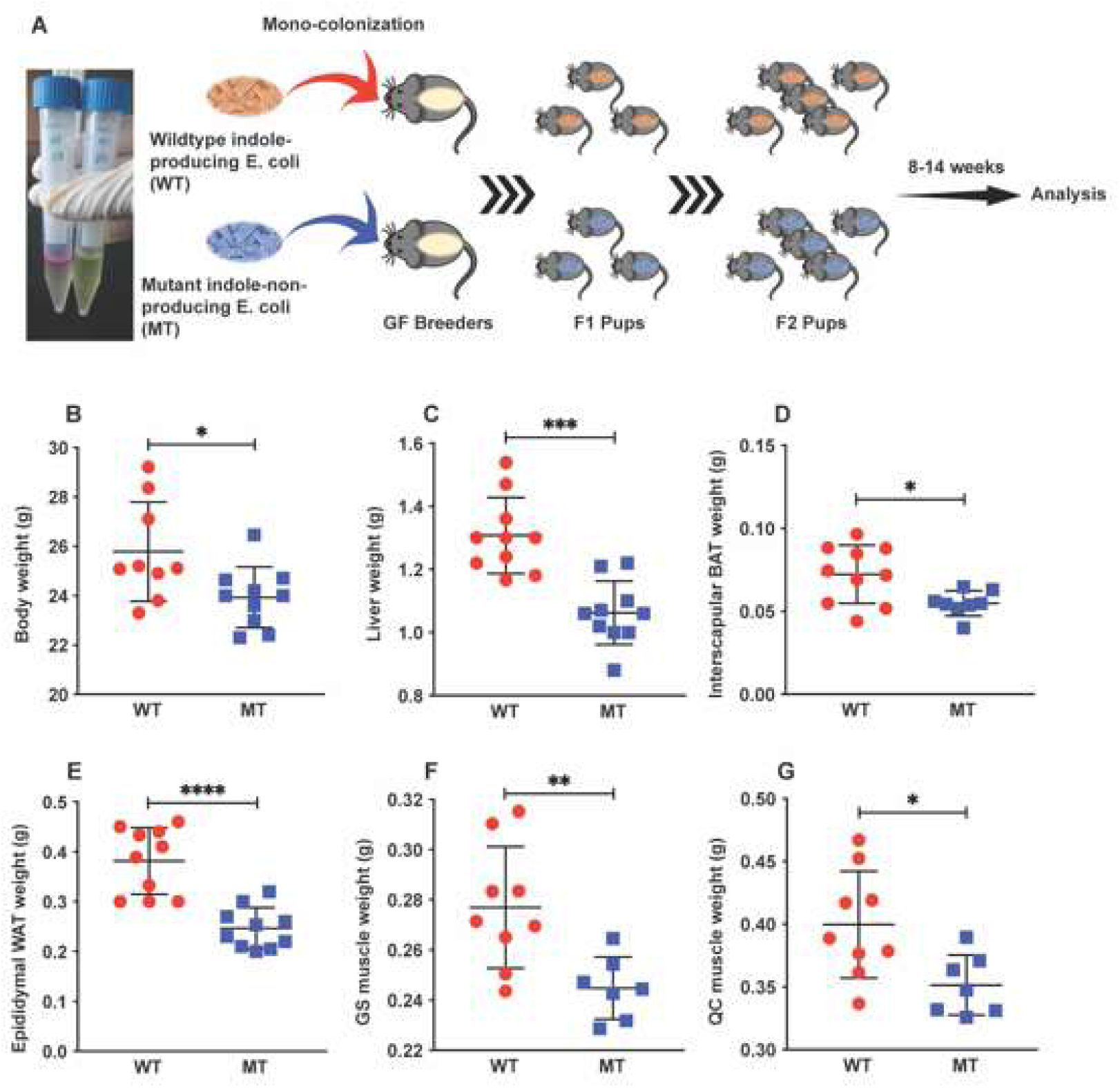
Germ-free (GF) mice colonized with indole-non-producing *E. coli* displayed lower body weights and lighter metabolic organs. **A** 6–8-week-old GF mice were colonized with wild type (indole-producing) and *tnaA*-mutant (indole-non-producing) *E. coli* (referred as WT and MT mice) by oral gavage. Second-generation (F2) pups were used for analysis to ensure their nursing by single gut microbial background parents. Photo shows Kovac indole tests on WT and MT *E. coli*, with the absence of the pink-colour indole ring in the MT sample. **B** Body weights of WT (n = 9) and MT (n = 10) mice at harvesting. **C** to **G** Weights of WT and MT mouse livers (C), interscapular brown adipose tissues (BAT) (D), epididymal white adipose tissues (WAT) (E), hindlimb muscles gastrocnemius (GS) (F), and quadriceps (QC) (G) (n = 7-10 per group). In B-D, data graphs show means ± SEM error bars. *p* values were calculated with the student’s *t* test. Statistical significance is presented as \**p* < 0.05, \*\**p* < 0.01, \*\*\**p* < 0.001, and \*\*\*\**p* < 0.0001 between indicated groups.

### Absence of gut microbial indole production alters serum metabolite profiles

Next, we performed targeted metabolite profiling using a combination of liquid chromatography-mass spectrometry (LC-MS) and nuclear magnetic resonance (NMR) methods on select metabolites involved in host metabolism, as well as host and gut microbial tryptophan metabolism. Compared to WT mice, MT mice had reduced levels of serum citrate and succinate (**Figure 2A, B**), two metabolic intermediates in the tricarboxylic acid (TCA) cycle. The TCA cycle converts acetyl-CoA from glycolysis to nicotinamide adenine dinucleotide (NADH) for oxidative phosphorylation and energy harvest. MT mice also had lower serum triglyceride and cholesterol levels (**Figure 2E, F**). These data, together with reduced expression of several key enzymes regulating the TCA cycle imply an impairment in mitochondrial metabolic activities, as both the TCA cycle and β-oxidation of fatty acids occur in the mitochondria. Interestingly, NAD^+^, the cofactor carrying electrons from glycolysis and the TCA cycle for ATP generation by oxidative phosphorylation in mitochondria, was present at lower, though not significantly different, levels in the serum of MT mice (**Figure 2C**). NAD^+^ levels are known to affect mitochondrial function, and reduced NAD^+^ levels may impair energy capture.

**Figure 2.**
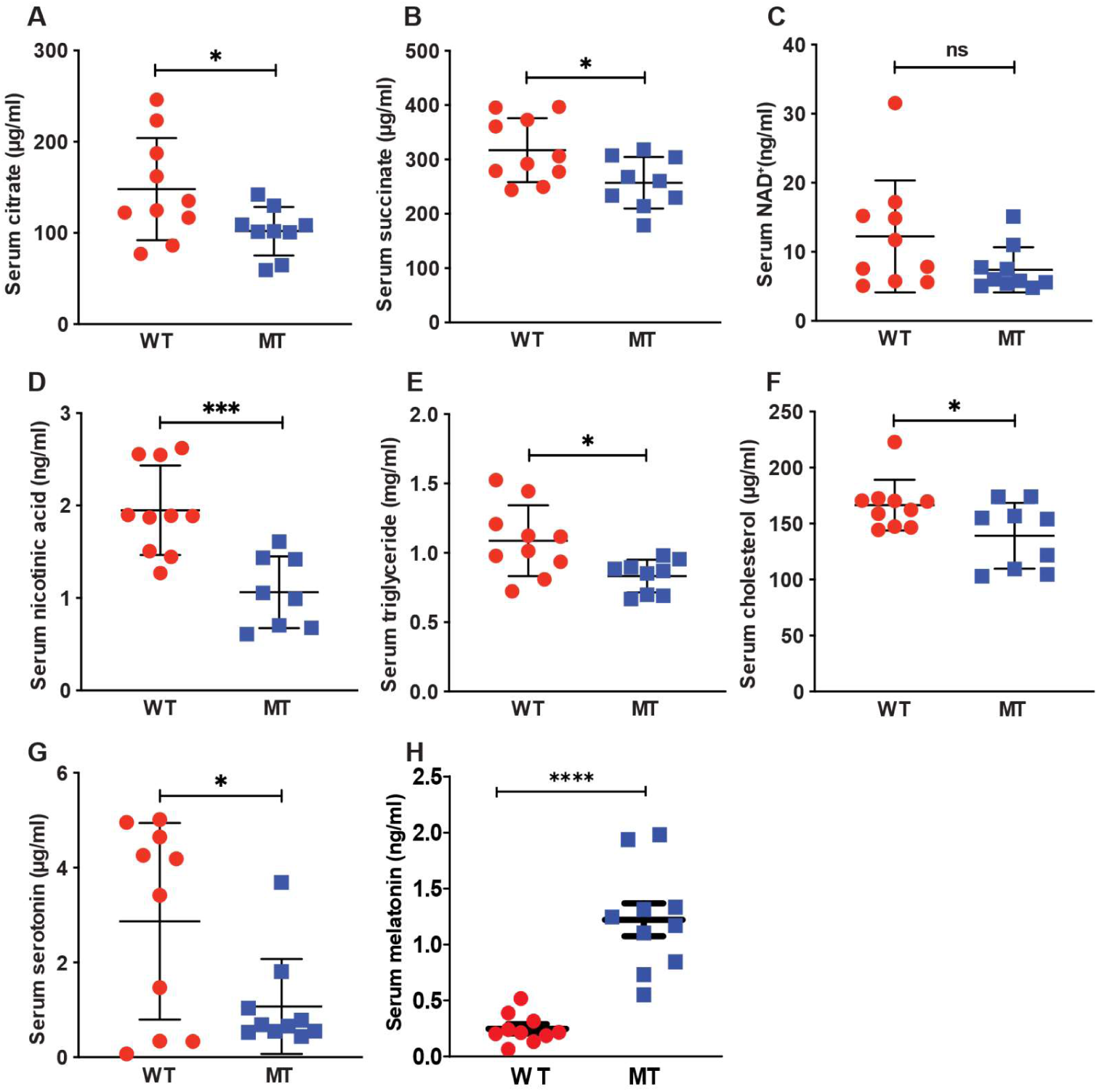
Loss of gut microbial indole production reduced concentrations of tricarboxylic acid (TCA) cycle intermediates, lipids, and serotonin in mouse serum. **A**, **B** Serum levels of TCA cycle intermediates citrate and succinate in WT (n=10) and MT (n=9) mice. **C** Serum level of nicotinamide adenine dinucleotide (NAD^+^) in WT and MT mice (n=10 per group). **D** Serum level of nicotinic acid (also known as niacin or vitamin B3) in WT (n=10) and MT (n=8) mice. **E, F** Serum levels of triglyceride and cholesterol in WT (n=10) and MT (n=9) mice. **G** Serum level of serotonin in WT and MT mice (n=10 per group). **H** Serum level of melatonin in WT and MT mice (n=10 per group). In A-H, data graphs show means ± S.D. *p* values were calculated with the student’s *t* test. Statistical significance is presented as \**p* < 0.05, \*\**p* < 0.01, and \*\*\**p* < 0.001 between indicated groups. ns denotes not significant with *p* ≥ 0.05.

In addition to microbial metabolism of tryptophan to produce indole and indole derivatives, the host also metabolizes tryptophan through the kynurenine-producing pathway and serotonin-producing pathway (**Supplemental Figure 2**) (Agus et al, 2018). The host kynurenine-producing pathway is upstream of host *de novo* NAD^+^ synthesis, contributing to the NAD^+^ pool (Agus et al, 2018). Apart from *de novo* NAD^+^ synthesis, the host may also produce NAD^+^ through salvage pathways, which utilize nicotinic acid (also known as niacin or vitamin B_3_) from dietary intake or gut microbial production. We examined the serum levels of key metabolites along these pathways (**Supplemental Table 1**) but found no significant difference in the serum level of tryptophan and, thus, its availability to the metabolic pathways. Most metabolites in the kynurenine-producing pathway did not differ significantly across the two groups. However, there was a significant reduction in the serum level of nicotinic acid in MT mice (**Figure 2D**), which may be a possible reason for the reduced serum NAD^+^ level (**Fig. 2C)** and a reduced serum level of the neurotransmitter serotonin (**Figure 2G**) and neuronal hormone melatonin (**Fig. 2H)**.

### Absence of gut microbial indole production alters host liver mitochondrial functions and lipid synthesis

As serum levels of key metabolites were reduced in MT mice, to gain deeper insight into the effect of microbial indole production on metabolic processes, we processed and analysed the main metabolic organs: liver, WAT, and muscles. The liver of MT mice contained lower levels of glycogen compared to that of WT mice (**Figure 3A**). We did not observe a significant difference in liver histology between the two groups (**Supplemental Figure 3**). However, hepatocyte nuclear factor-4α (*Hnf4a*), a regulator of genes involved in glucose transport and glycolysis, was expressed at slightly lower levels in the livers of MT mice than WT mice (**Figure 3B**). Expression levels of enzymes involved in the TCA cycle (*Pc*, *Cs*, *Aco2*) and lipogenesis enzymes (*Acly*, *Acaca*, *Acacb*) were also reduced in the MT mouse liver (**Figure 3B, C**), which correlates with the reduced serum levels of citrate and succinate (**Figure 2A, B**).

**Figure 3.**
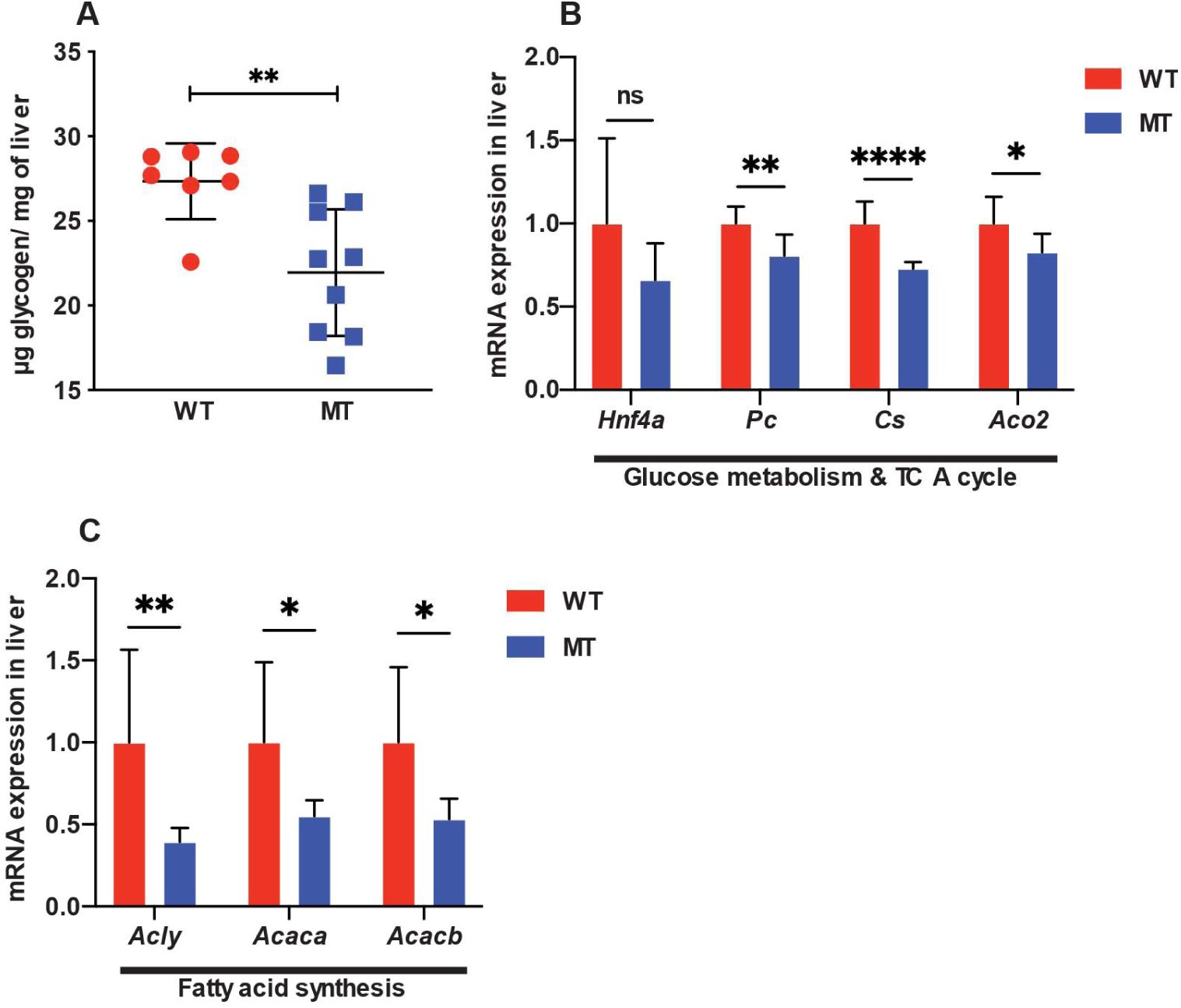
Indole-non-producing *E. coli*-colonized mice show signs of reduced glucose metabolism and lipogenesis in livers. **A** Liver glycogen levels (μg glycogen per mg tissue) (WT n=7, MT n=9). **B** mRNA expression of glucose metabolism regulator *Hnf4a*, TCA cycle enzymes *Pc* (pyruvate carboxylase), *Cs* (citrate synthase), and *Aco2* (aconitase 2) in the liver tissues of WT and MT mice (n=8 per group). **C** mRNA expression of ATP citrate lyase (*Acly*, converting citrate to acetyl-CoA), and acetyl-CoA carboxylase (*Acaca* and *Acacb*, converting acetyl-CoA to malonyl-CoA, the rate-limiting step in fatty acid synthesis) in the livers of WT and MT mice (n=7-8 per group). mRNA levels were normalized to the control gene *Hprt1*. In A-C, data graphs show means ± S.D. *p* values were calculated with the student’s *t* test. Statistical significance is presented as \**p* < 0.05, \*\**p* < 0.01, \*\*\**p* < 0.001, and \*\*\*\**p* < 0.0001 between indicated groups. ns denotes not significant with *p* ≥ 0.05.

Interestingly, in the epididymal WAT of MT mice, the mRNA levels of glyceroneogenesis rate-limiting enzyme *Pck1*, and another glyceroneogenesis enzyme *Pck2*, were upregulated (**Figure 4B**). Glyceroneogenesis, which generates glycerol 3-phosphate from precursors other than glucose, such as pyruvate, typically occurs during fasting or starvation in WAT. Production of glycerol 3-phosphate enables the re-esterification of free fatty acids in adipose tissue, which is a concomitant process with active lipolysis (Hanson & Reshof, 2003; Leithner et al, 2018). Consistently, the mRNA levels of regulators of lipolysis *Adrb2/3*, as well as long-chain fatty-acid-coenzyme A ligase *Acsl1*, were significantly upregulated (**Figure 4C**). Coupled with the upregulation of lipolysis genes was the decreased activation of cyclic AMP-response element binding protein (CREB), a primary regulator of lipogenesis in the WAT of MT mice (**Figure 4A**). Lipogenesis enzyme mRNA expression was also downregulated in the WAT (i.e., *Scd2*, **Figure 4C**) of MT mice. Corresponding to the reduced weight of WAT in MT mice (**Figure 1E**), morphological analysis of the adipocytes showed more cells with smaller areas (**Supplemental Figure 4C**) and slightly higher total cell number per selected area (**Supplemental Figure 4B**). These results suggest that, compared to WT mice, MT mice have elevated lipolysis in the WAT, indicative of metabolic stress.

**Figure 4.**
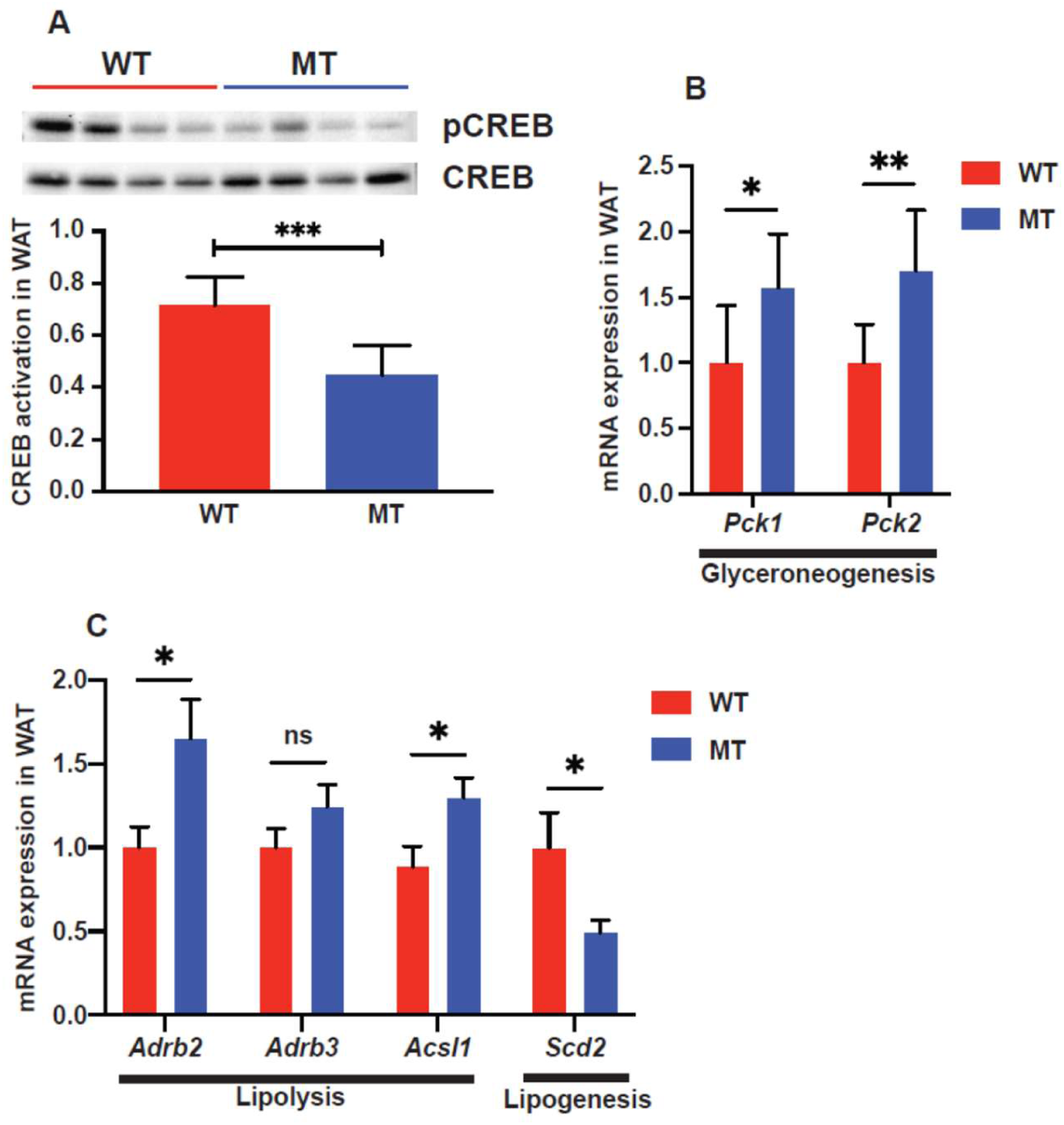
White adipose tissues (WAT) of indole-non-producing *E. coli*-colonized mice showed elevated glyceroneogenesis and lipolysis. **A** Western blotting and quantification of activation of cyclic AMP-response element binding protein (CREB), a lipogenesis regulator, in epididymal WAT of WT and MT mice (n=6-8 per group). Phosphorylated CREB (pCREB), a measure of CREB activation, was normalized to total CREB. **B** mRNA expression of *Pck1* and *Pck2* (phosphoenolpyruvate carboxykinase 1/2, cytosolic/mitochondrial, respectively) in epididymal WAT of WT and MT mice (n=7-8 per group). Phosphoenolpyruvate carboxykinases catalyse a rate-limiting step in glyceroneogenesis in WAT, the process which generates glycerol 3-phosphate from precursors other than glucose typically during fasting. **C** mRNA expression of lipolysis regulators *Adrb2/3* (β_2/3_ adrenoreceptor), long-chain fatty-acid-coenzyme A ligase *Acsl1*, and fatty acid synthesis enzymes stearoyl-CoA desaturase (*Scd2*) in epididymal WAT of WT and MT mice (n=6-8 per group). mRNA levels were normalized to the control gene *Hprt1*. In A-C, data graphs show means ± S.D. *p* values were calculated with the student’s *t* test. Statistical significance is presented as \**p* < 0.05, \*\**p* < 0.01, and \*\*\**p* < 0.001 between indicated groups. ns denotes not significant with *p* ≥ 0.05.

Similar to the liver, the QC muscles contained lower levels of glycogen compared to that of WT mice (**Figure 35A**). Glucose metabolism and mitochondrial functional genes [e.g., *Glut4* (a glucose transporter), succinate dehydrogenase (*Sdh*; a TCA cycle and respiratory chain enzyme), and *Tfam* (a key activator of mitochondrial transcription)] were downregulated in the MT mouse GS muscles compared to WT muscles (**Figure 5B**). Corresponding to the reduced glucose metabolism in the GS muscles of MT mice, the mRNA expression of myosin heavy-chain II proteins *Myhc-2b* and *Myhc-2d*, the “fast” isoforms of myosin heavy-chain proteins (i.e., utilize glycolysis), were downregulated, along with *Rapsyn*, a neuromuscular junction-associated gene (**Supplemental Figure 5**). Interestingly, muscle atrophy-related genes *Murf-1* and *Atrogin-1* (encoding E3 ubiquitin ligases) and myogenesis-related gene *Myod* were downregulated in the GS muscles of MT mice (**Supplemental Figure 5**).

**Figure 5.**
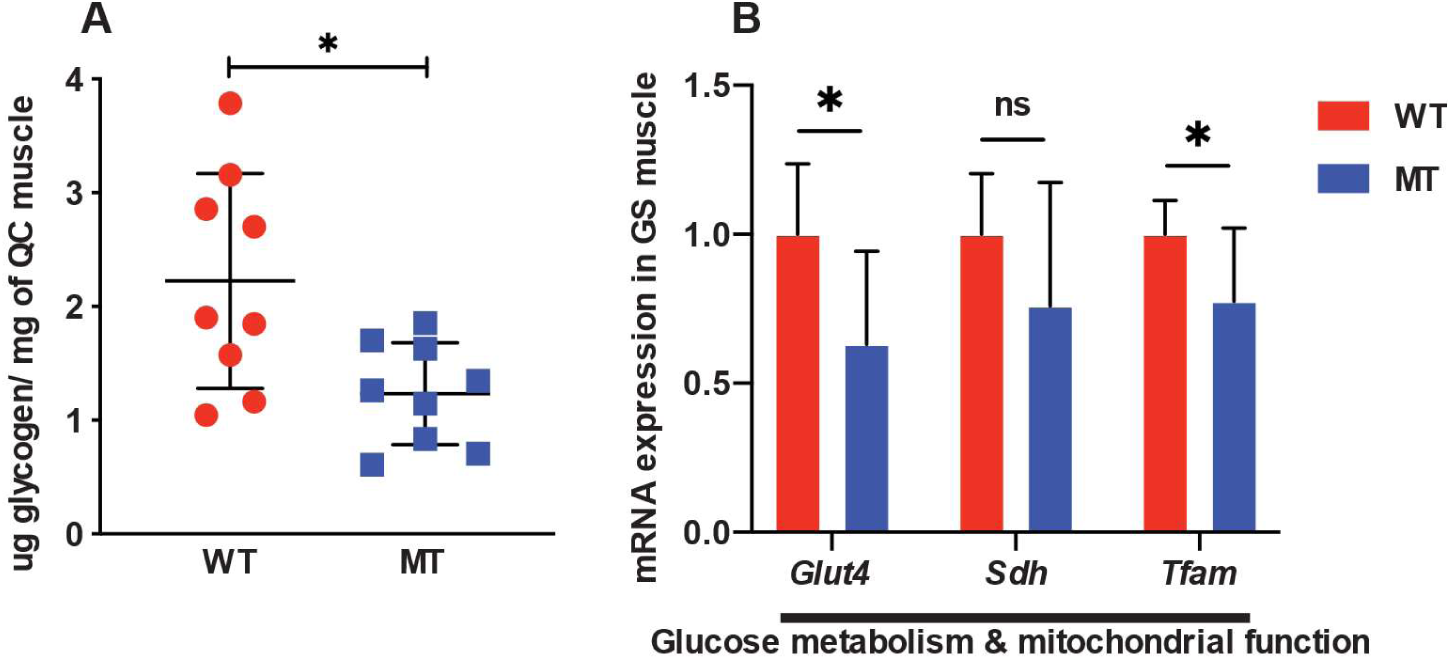
Indole-non-producing *E. coli*-colonized mice contained reduced glycogen level in hindlimb muscles. **A** Glycogen levels in hindlimb muscle quadriceps (QC) (μg glycogen per mg tissue) (WT n=8, MT n=9). **B** mRNA expression of glucose transporter *Glut4*, TCA cycle and respiratory chain enzyme succinate dehydrogenase *Sdh*, and a key activator of mitochondrial transcription *Tfam* (mitochondrial transcription factor A), in hindlimb gastrocnemius (GS) muscle tissues of WT and MT mice (n=7-8 per group). In A-B, data graphs show means ± S.D. *p* values were calculated with the student’s *t* test. Statistical significance is presented as \**p* < 0.05, \*\**p* < 0.01, and \*\*\**p* < 0.001 between indicated groups. ns denotes not significant with *p* ≥ 0.05

### MT mice exhibit increased food consumption without elevated energy expenditure

The low body weight, reduced serum levels of TCA cycle intermediates and lipids, and lower liver and muscle glycogen levels combined with elevated lipolysis and reduced lipogenesis gene expression in the liver and adipose tissues suggest a phenotype of metabolic stress in MT mice. There are three possible explanations for this observation: reduced food consumption, reduced intestinal function to digest and absorb nutrients, or increased metabolic rate (spending more energy). Therefore, we performed metabolic cage experiments to monitor food and water intake, oxygen consumption, carbon dioxide production, heat production, and locomotion at room temperature (22°C).

MT mice had similar food and water intake at night compared to WT mice but exhibited significantly elevated food consumption during the daytime (**Figure 6A, B**). However, the respiratory exchange rate, heat production, and calculated daily energy expenditure did not differ between the two groups (**Figure 6C** to **E**). Notably, there was an increase in the locomotor activity in the MT mouse group (**Figure 6F**, **G**), though the increased activity did not reflect the daily energy expenditure. Calorimetric cage analyses at cold (16°C) temperature (**Supplemental Figure6**) showed a similar respiratory exchange rate, heat production, and daily energy expenditure. However, at a thermoneutral temperature (30°C), at which the mammals maintain the minimum metabolic rate (basal metabolic rate) (Skop et al, 2020), MT mice had a lower respiratory exchange rate and slightly higher heat production and daily energy expenditure (**Supplemental Figure6C** to **E**). As the respiratory exchange rate is indicative of the sources of metabolic fuels (carbohydrate, protein, or fat) utilized, the lower respiratory exchange ratio could indicate higher lipid catabolism at basal metabolism. Taken together, these data do not explain the phenotypic observations in the MT mice.

**Figure 6.**
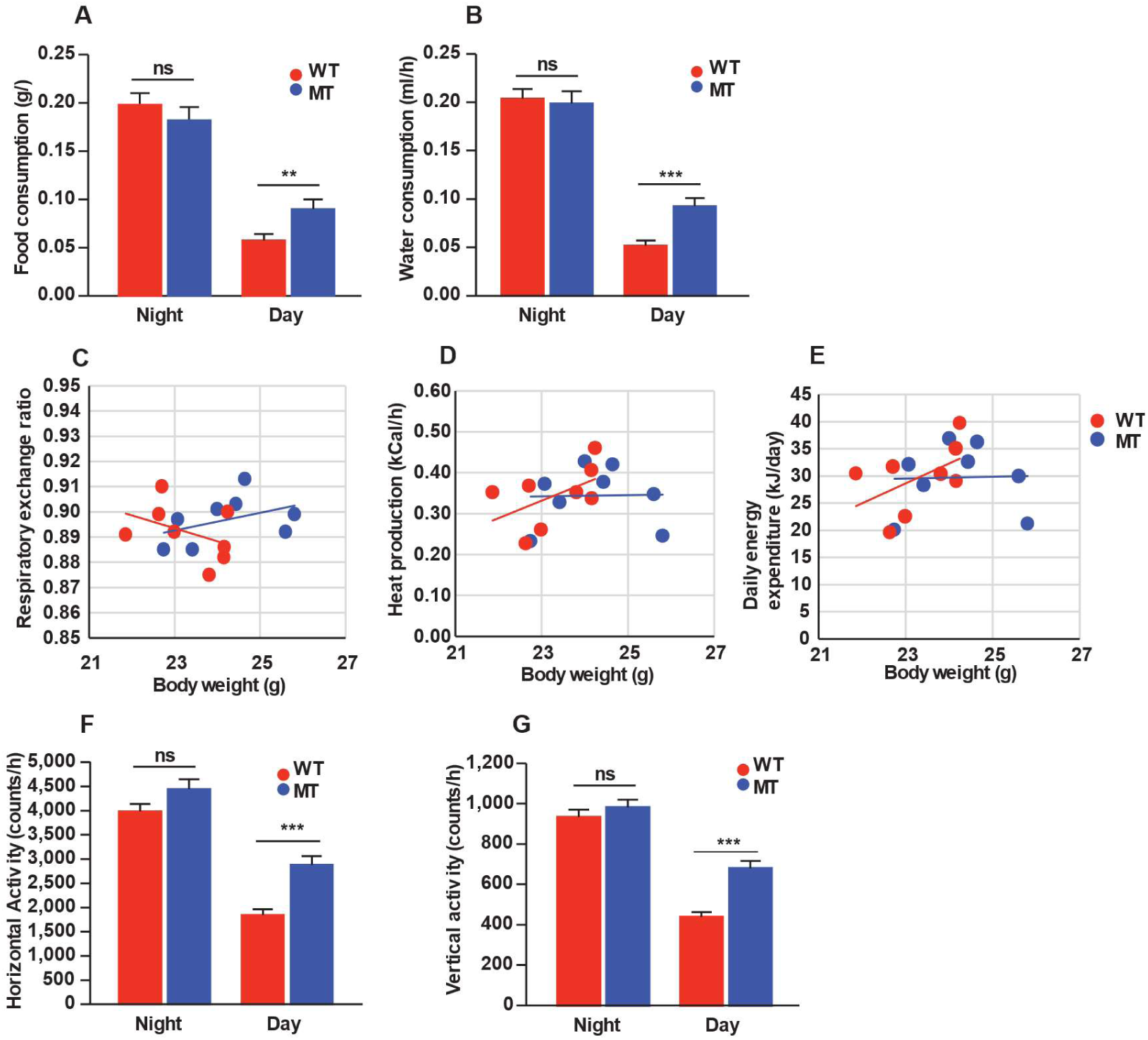
Mice with indole-non-producing *E. coli* showed increased food intake and locomotor activity through calorimetric cage analysis. **A**, **B** Food and water consumption at night- and daytime of WT and MT mice. **C** to **E** Linear regression curves for correlation of body weight on the respiratory exchange rates (C), heat production (D), and daily energy expenditure (E). Data are the means of results over three experimental days per animal. Each dot represents an animal. **F**, **G** Horizontal and vertical activities of WT and MT mice. Experiments were conducted at a constant temperature of 22°C and humidity of 67%. A total of 3 days in the calorimetric cages were used for the data analysis. n=8 per group. For **A**, **B**, **F**, **G**, data graphs show means ± SEM error bars. *p* values were calculated with ANOVA (Welch’s robust *t* test of equality of means). Statistical significance is presented as \**p* < 0.05, \*\**p* < 0.01, \*\*\**p* < 0.001, and \*\*\*\**p* < 0.0001 between indicated groups. ns denotes not significant with *p* ≥ 0.05. For **C-E**, data were analysed by ANCOVA GLM statistical analysis comparing the two groups with the body weight and day/night variation as covariates.

### MT mice have increased intestinal length, enlarged caecum, increased epithelial cell growth and fewer serotonin-producing colonic enterochromaffin cells

The results thus far that MT mice were eating more and not burning more energy but still had reduced body weight/wasting syndrome phenotype suggested a possible digestion and or absorption problem in the intestinal tract. Indeed, MT mice had a longer small intestine (SI) and colon (**Figure 7A** to **C**) and an enlarged caecum filled with semi-digested food (**Figure 7D**). Further histological analysis of the jejunal epithelium showed no difference in the villus length and number (**Figure 7E** to **G**), but significant changes were observed in the crypt depths (**Fig 7H**), consistent with a possible compensatory mechanism to increase the absorption area in MT mice.

**Figure 7.**
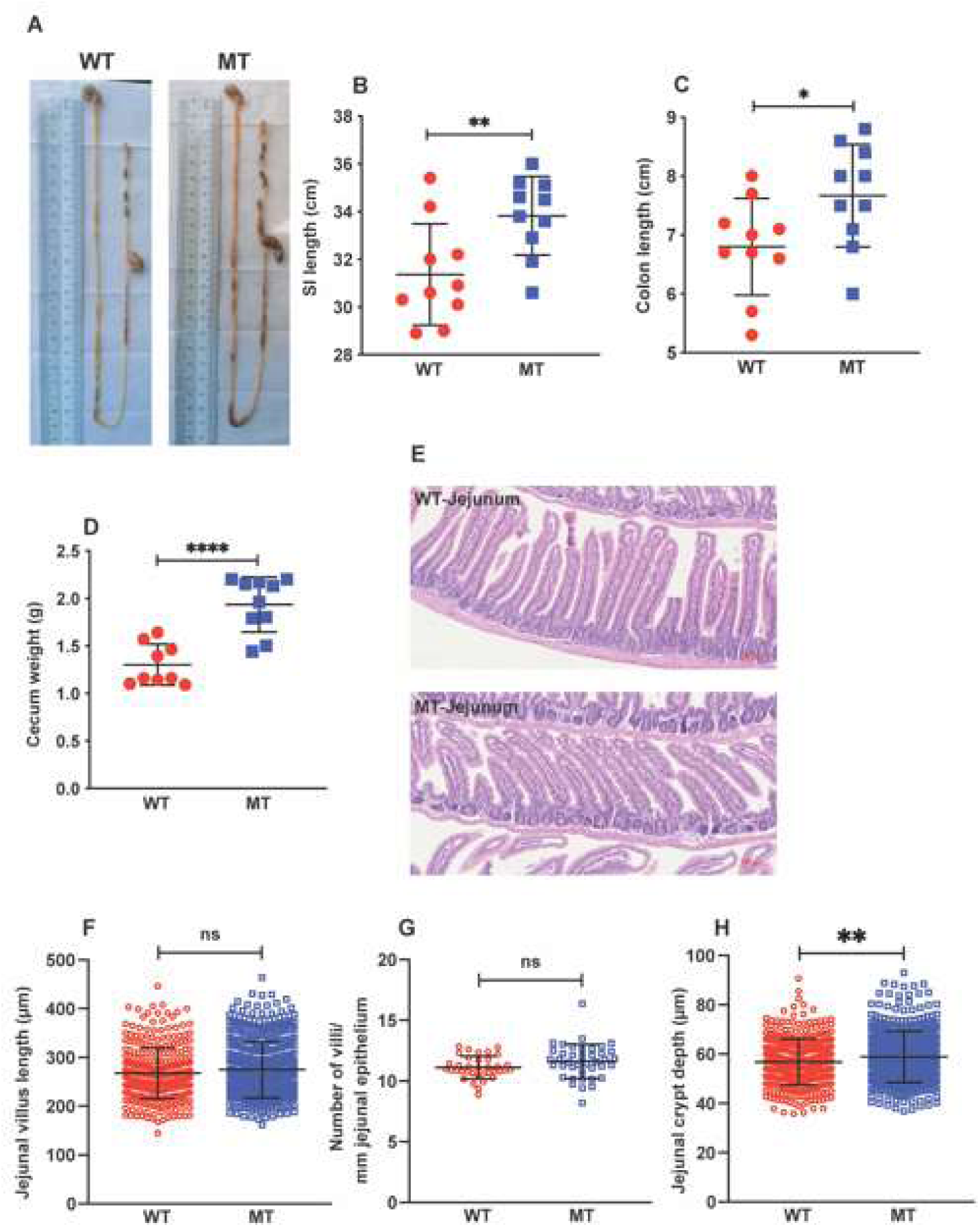
Mice with indole-non-producing *E. coli* showed enlarged caecum without drastic changes in general histology. **A** Representative image of gastrointestinal tracts of WT and MT mice when harvested. **B** to **D** Length of small intestines (SI, B) and colons (C), and weight of caeca (D) of WT and MT mice (n=9-10). **E** Representative images of haematoxylin and eosin (H&E) staining of jejuna of WT and MT mice. Scale bar, 100 μm. **F** to **H** Quantification of villus length (F), villus number (G), and crypt depth (H) in the jejuna of WT and MT mice (n=6 per group). In B-D and F-H, data graphs show means ± S.D. *p* values were calculated with the student’s *t* test. Statistical significance is presented as \**p* < 0.05, \*\**p* < 0.01, \*\*\**p* < 0.001, and \*\*\*\**p* < 0.0001 between indicated groups. ns denotes not significant with *p* ≥ 0.05.

The intestinal barrier integrity and inflammatory status also play crucial roles in normal digestive and absorptive functions (Vancamelbeke & Vermeire, 2017). Therefore, we screened a panel of barrier integrity and inflammation markers in the SI epithelium. We found no differences in gene expression of the tight junction proteins *Occludin*, *Zo-1*, and *Claudin-1*, or the pro-inflammatory cytokines *Tnf*a, *Il1b*, and *Il*18 between the two groups (**Supplemental Figure7**).

Having observed a reduced level of circulating serotonin in MT mice (**Figure 2G**), we quantified enterochromaffin (chromogranin A-expressing, CHGA^+^) cells by immunohistochemistry in the SI and colon. We found a significant reduction in the number of CHGA^+^ cells in the colon, but not in the SI of MT mice (**Figure 8A, B**; **Supplemental Fig. 8A, B**), consistent with reduced gut motility and digestive function as observed in mice with reduced serotonin levels (Ge et al, 2017). Gene expression analysis of the enzymes involved in serotonin synthesis and degradation in the SI and liver showed similar results between the two groups (**Supplemental Figure8C, D**).

**Figure 8.**
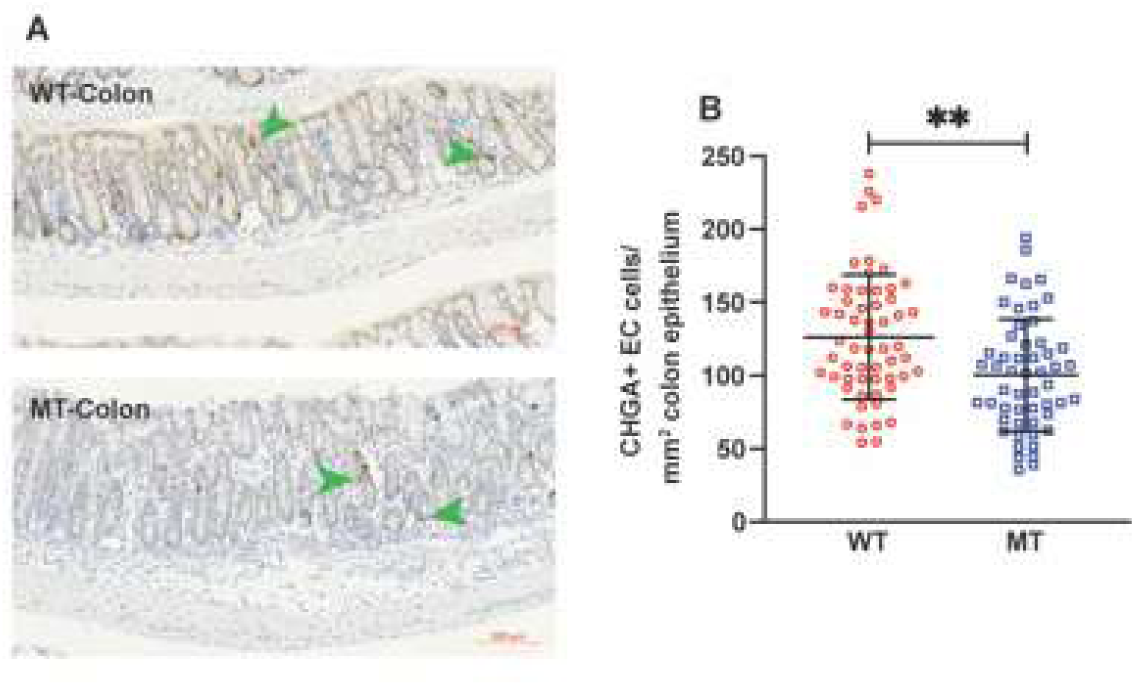
Indole-non-producing *E. coli*-colonized mice showed reduced colonic serotonin-producing enterochromaffin cells. **A** Representative images of colonic tissue sections stained with anti-chromogranin A (CHGA) antibody for enterochromaffin cells (in brown, indicated by green arrow heads) of WT and MT mice. Nuclei were counterstained with haematoxylin (in blue). Scale bar, 100 μm. **B** Quantification of CHGA^+^ enterochromaffin cells (number of cells per mm^2^ colonic epithelium) in the colons of WT and MT mice (n=6 per group). In A-B, data graphs show means ± S.D. *p* values were calculated with the student’s *t* test. Statistical significance is presented as \*\**p* < 0.01 between indicated groups.

Mucus-secreting goblet cells are another major group of secretory cells in the intestinal epithelium. We performed periodic acid-Schiff (PAS) staining on the intestines and found a decrease in the number of goblet cells in the jejunal villi of MT mice (**Supplemental Supplemental Figure 9A, B**). However, no change in the mRNA expression of *Muc2* was observed in the MT group (**Supplemental Supplemental Figure 9C**).

Previous histological analysis revealed deeper intestinal crypts in the jejuna of MT mice (**Figure 7H**). As intestinal crypts contain proliferating stem cells and transit amplifying progenitor cells (Gehart & Clevers, 2019), we performed immunohistochemical staining for cellular proliferation marker KI67^+^, which confirmed a marked increase in the number of proliferating KI67^+^ cells in the SI of MT mice (**Figure9A, B**). As indoles are known to elicit cell cycle exit (Chinni et al, 2001), our findings in the MT mice may not be surprising. In addition, we noticed a reduced expression of the intestinal active stem cell marker *Lgr5* (**Fig. 9C**) consistent with continuous signals from the mTOR pathway, which is known to push stem cell towards cell differentiation, thus reducing the recycling and renewal of *Lgr5* positive progenitor cells (Li et al, 2021).

**Figure 9.**
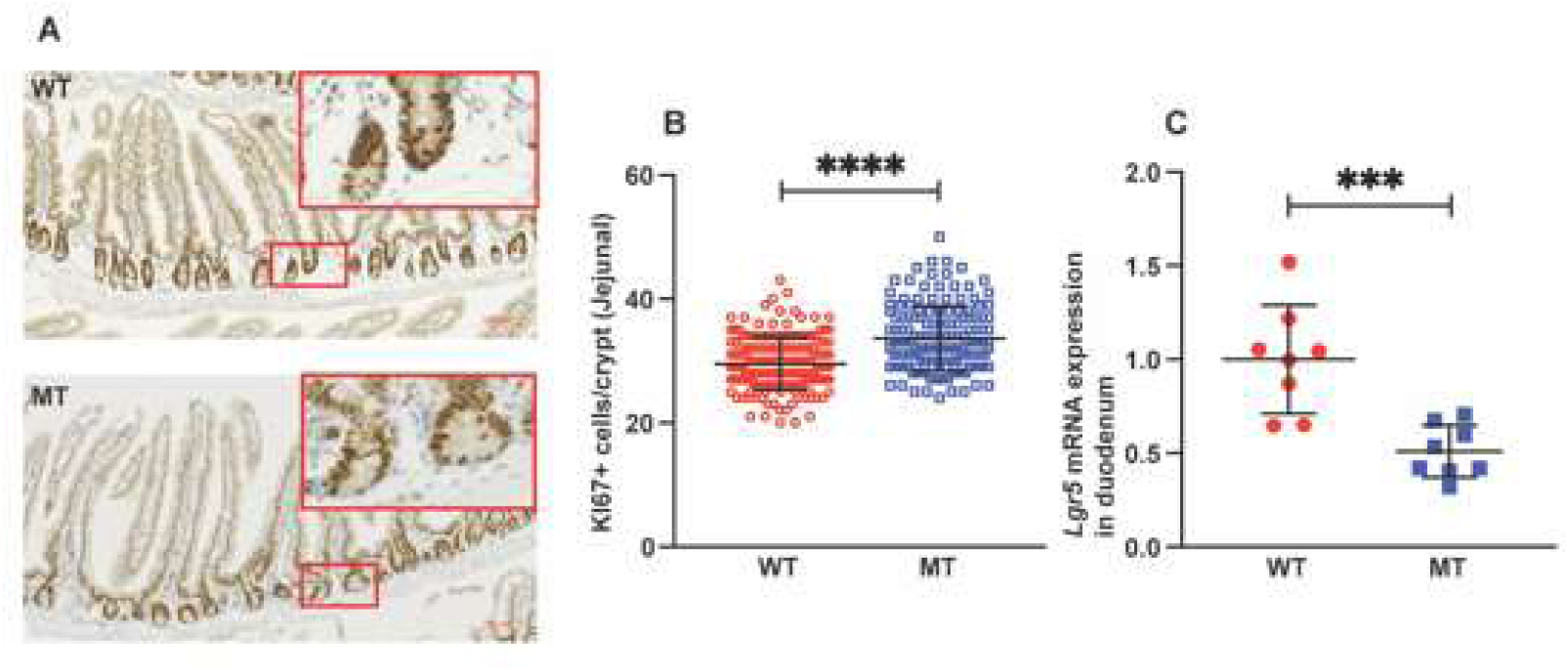
Mice colonized with indole-non-producing *E. coli* showed elevated SI epithelial growth. **A** Representative images of jejunal tissue sections stained with anti-KI67 antibody for proliferating cells (brown), in jejuna of WT and MT mice, with haematoxylin-counterstained nuclei (blue). Scale bar, 100 μm. **B** Quantification of KI67-expressing (KI67^+^) proliferating cells (number of cells per crypt) in the jejuna of WT and MT mice (n=6 per group). **C** mRNA expression of intestinal active stem cell marker *Lgr5* in the duodena of WT and MT mice (n=8 per group). mRNA levels were normalized to the control gene *Hprt1*. In A-C, data graphs show means ± S.D. *p* values were calculated with the student’s *t* test. Statistical significance is presented as \*\*\**p* < 0.001 and \*\*\*\**p* < 0.0001 between indicated groups.

### MT mice have lower mRNA expression of *kcnj12* in the liver and small intestine

One mechanism accounting for the prolonged health span phenotype is the indole-aryl hydrocarbon receptor (AhR) signalling pathway (Sonowal et al, 2017). The ligand activation of AhR is also known to regulate *kcnj12* (Obata et al, 2020), the gene encoding an inwardly rectifying K+ channel with relevance to gut motility by establishing action potential waveform and excitability of neuronal tissues and is known to regulate mitochondrial membrane potential and mitochondrial ROS production (Lee et al, 2010; Obata et al, 2020). In exploring a possible mechanism underlying the observed changes in intestinal motility and liver function in MT mice, we assessed the expression of *kcnj12.* Both the liver and intestine of MT mice had considerably reduced expression of *kcnj12* (**Fig. 10)**. These findings provide additional explanations for the reduced gut motility due to impairment in the SI smooth muscle function and mitochondrial function in the liver.

**Figure 10.**
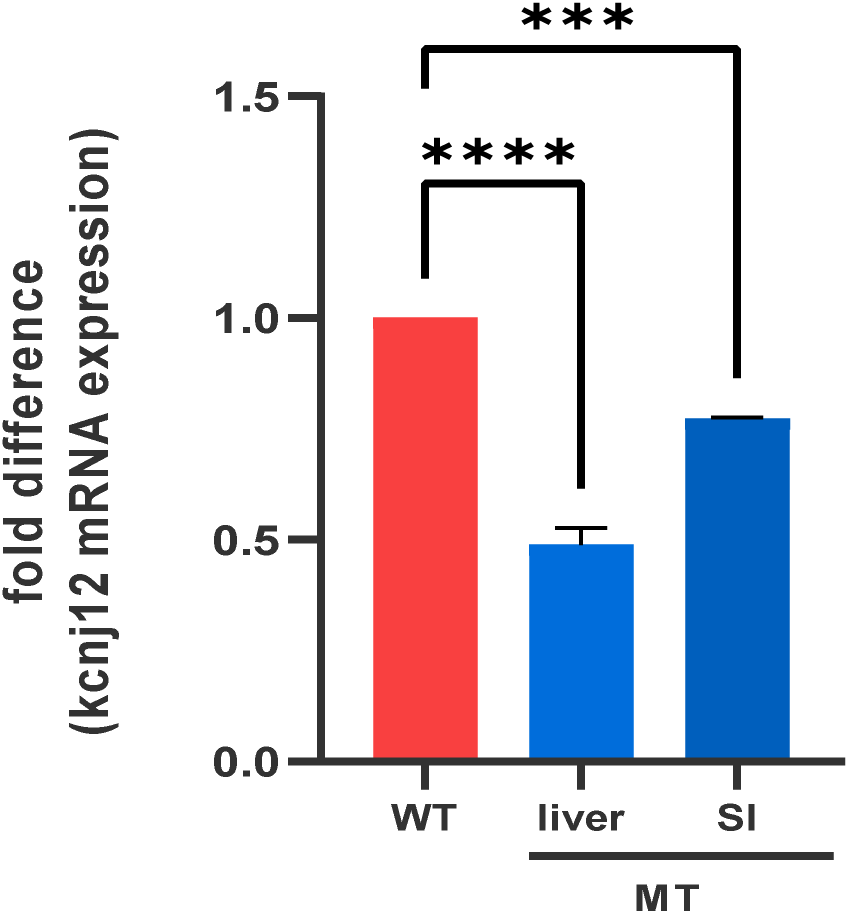
Mice colonized with indole-non-producing *E. coli* showed reduced levels of *Kcnj12* mRNA expression in the liver and small intestine. Fold difference in the mRNA expression of *Kcnj12* in the liver and small intestine (SI) of WT and MT mice (n=8 per group). mRNA levels were normalized to the control gene *Hprt1*. Data graphs show means ± SEM. *p* values were calculated with One-way ANOVA. Statistical significance is presented as \*\**p* < 0.01 and \*\*\*\**p* < 0.0001 between indicated groups. ns denotes not significant with *p* ≥ 0.05.

## DISCUSSION

Here, we report a wasting syndrome phenotype using a genetically intact and non-manipulated GF mouse colonized with microbes unable to produce indoles. The data connects microbe derived indoles to the regulation of host metabolism as observed by multiple organ decline associated with impairment in the TCA cycle, reduced levels of triglycerides and cholesterol, dysfunctional glucose metabolism in the liver, lipolysis, and reduced white adipocyte tissue, liver weight and skeletal muscle mass. Calorimetric chamber experiments revealed that indole mutant mice show increased food intake but comparable daily energy expenditure. Detailed profiling of the intestinal canal revealed large quantities of undigested food resulting in increased caecum pool in MT mice. Further profiling revealed reduced numbers of enterochromaffin cells resulting in reduced levels of circulating serotonin. Collectively these findings are consistent with major impairment in the digestive function of MT mice due to reduced gut motility. The enlargement of the caecum in MT mice resembles the characteristic phenotypes displayed by GF and antibiotic-treated rodents, in which enlarged caeca are filled with undigested material (Wostmann, 1981). Caecal enlargement induced by antibiotic treatment in mice is associated with reduced intestinal motility, impaired digestion, and longer gut transit time, which ultimately impacts energy harvesting and body weight (Wostmann, 1981; Ge et al, 2017). These changes are often observed in ageing mice and imply a link between organ decline and function in the ageing body (Soenen et al, 2016). Furthermore, previous studies have established the role of gut microbiota in the modulation of colonic enterochromaffin cell numbers and serotonin production and, thus, their role in gut motility and digestion (Yano et al, 2015; Ge et al, 2017). In addition, the observation of a longer intestine and increased epithelial growth in MT mice, are consistent with a possible compensatory mechanism to increase the absorption surface and capacity (Chevalier et al, 2015). Interestingly, depletion of gut microbiota by antibiotic treatment caused mice to have increased SI length, like the phenotype of GF mice (Chevalier et al, 2017). The alterations in the metabolomic profiles and data suggestive of malnourishment in the MT mice support this possibility. However, further investigations are needed to verify this speculation.

Mice treated with a broad-spectrum antibiotic cocktail present with lower colonic serotonin levels, reduced colonic motility, and phasic contractions of the longitudinal colonic muscles (Ge et al, 2017). On the other hand, mono-colonization of GF mice with *Clostridium ramosum* succeeds in promoting enterochromaffin cell differentiation and serotonin production, facilitating intestinal fatty acid absorption and modulating metabolic functions (Mandic et al, 2019). The MT mice presented reduced serum serotonin and fewer colonic enterochromaffin cells consistent with a gut microbe-indole-mediated feed-forward mechanism to support serotonin production in the colon, therefore tuning gut motility, digestion, and host metabolic homeostasis. A recent report using mice with enteric neuron-specific deletion of AhR showed that the lack of AhR in these neurons resulted in significantly prolonged intestinal transit time (ITT) (Obata et al, 2020). This observation from Pachnis team is directly in line with our results in MT mice suggesting reduced gut motility, as indoles are ligands for AhR activation (Hubbard et al, 2015). Transcriptomics analysis of enteric neurons for downstream AhR targets that were also potential regulators of neuronal activity in these neurons identified *kcnj12*, a gene that encodes the inwardly rectifying K+ channel, subfamily J member 12 (Kir2.2) (Obata et al, 2020), which is known to regulate the excitability of cardiac muscles and neurons (Hibino et al, 2010). MT mice had a significant decrease in the expression of *Kcnj12* in the liver and SI consistent with reduced indole-AhR signalling. It is tempting to speculate that this reduction in *kcnj12* expression in the SI and livers of MT mice contributes, in part, to lower neuronal excitability and, thus, reduced gut motility, and metabolic impairment in the mitochondria function as knock-down of *Kcnj12* has been shown to significantly increase ROS activation, trigger cell cycle arrest and mitochondria dysfunction (Lee et al, 2010).

In addition to the reduction in serum levels of the TCA cycle intermediates and lipids, altered concentrations of other metabolites were found in MT mice. Many of the biomarkers observed in this study are associated with increased ROS activity and/or impairment in mitochondrial function consistent with the reduced expression of Kcnj12. Additional enzymes include citrate synthase, the first enzyme in the TCA cycle and a known quantitative indicator of intact mitochondria (Scaini et al, 2010), leading to a reduction in citrate in MT mice (**Fig. 2A)**. Melatonin is well-known for its anti-oxidative stress effect (Galano & Reiter, 2018), and the significant increase in serum melatonin in MT mice (**Fig 2H and Supplemental table 1**) further suggests elevated levels of ROS activity in indole-deficient MT mice. Interestingly, knock-down of Kcnj12 activity has been shown to significantly increase ROS activation, triggering cell cycle arrest and mitochondrial dysfunction (Lee et al, 2010). Indoles are potent radical scavengers that protect against oxidative stress (Dodd et al, 2017). Given that reactive oxygen species (ROS) production typically increases with age, maintaining indole concentrations as one gets older may be beneficial to reduce age-associated ROS levels and age-related metabolic disorders (Furukawa et al, 2004). Though we have not specifically addressed the details of mitochondrial impairment and ROS production in this study, it is tempting to speculate a possible link between the indole-AhR signalling pathway and regulation of mitochondrial function and ROS homeostasis, which warrants further investigation.

Another interesting observation is the increase in picolinic acid (PA) (**Supplemental table 1**), an endogenous metabolite of tryptophan known to have a beneficial effect by converting tryptophan to nicotinamide (Shibata & Fukuwatari, 2014). Our metabolomic analysis show reduced levels of NAD^+^, though not statistically significant (**Supplemental table 1**). However, the increased levels of PA may be a compensatory mechanism imposed in the indole-mutant mice.

Chronological age is an important predictor of morbidity and mortality but is insufficient to account for the individual variation in declining organ function with advancing age. Therefore, the idea of developing a metabolomic signature(s) of biological aging that could predict changes in organ function is attractive and was recently applied to better define the biological age among old human people. Among the metabolites associated with a younger biological age in humans is the microbial metabolite indole-3 acetic acid (IAA) (Johnson et al, 2019). Notably, the indole mutant mice display massive reduction in plasma levels of IAA (**Supplemental table 1**). While further experiments are warranted, these findings suggest that the MT mice are experiencing signs of accelerated biological ageing thus providing clues into the underlying molecular mechanisms that regulate biological ageing through interactions between the gut microbe metabolites and host organs. The previous observation that prolonged exposure to indoles in flies, worms and mice associated with prolonged health span support the findings presented here (Sonowal et al, 2017). It is obvious that more detailed analyses are required to reveal the full scope of physiological and biochemical changes in the indole mutant mice. A detailed hormone profiling has not been performed nor are the ROS-mitochondria dysfunctions dissected in detail. Yet, this report underscores the importance of exploring gut microbiota–host interactions at systems biological level if we aim to identify individuals at risk of accelerated biological ageing. Our reductionist approach highlights the gut microbe derived tryptophan metabolites, indoles and I3A, to be interesting representatives of the next generation set of biomarkers to be included to the 21^st^ century precision medicine healthcare repertoire to prolong human health span.

## MATERIALS AND METHODS

### Animal Experiments

All animal works were carried out with the approval of the Institutional Animal Care and Use Committees (IACUC) of the Nanyang Technological University (NTU), Singapore (Reference numbers: AUP-E0016 & AUP-A19104) and SingHealth, Singapore (Reference number: 2016/SHS/1263), and in accordance with the guidelines of the Responsible Care and Use of Laboratory Animals (RCULA) set out by the National Advisory Committee for Laboratory Animal Research (NACLAR) in Singapore. All mice were housed in designated rooms at the NTU Animal Research Facility (NTU-ARF) and the SingHealth Experimental Medicine Centre (SEMC), with a 12 h light-dark cycle and *ad libitum* food and water intake, unless otherwise specified.

Germ-free (GF) C57BL/6J mice aged 6 to 8 weeks were colonized with wild type (indole-producing) or *tnaA* mutant (indole-non-producing) *Escherichia coli*. Based on the weight of the mouse, 100 to 250 μL of the bacteria suspension was given to each mouse by oral gavage. After gavage, a maximum of 5 mice were housed in GF bubble tents. Mice with wild type *E. coli* were separated from mice with mutant *E. coli*. Forty-eight hours, 1 week, and 2 weeks following the gavage, mouse body weights were recorded, and stool samples were collected. Suspension of stool samples was used for streaking on bacterial culture agar plates to examine colonization and contamination. Once the colonization in the mice was confirmed, male and female mice were paired at a ratio of 1:2 at 8 weeks old for breeding. Second generation litters were used for analysis. Pups were weaned at 3 weeks old and monitored for their growth characteristics. At the end points (8-14 weeks), their guts, livers, muscles, adipose tissues, brains, and blood were harvested for further processing and analysis.

### Bacterial culture

Wild type and *tnaA* (encoding tryptophanase) mutant *Escherichia coli* strains K12 were kind gifts from Dr Thomas K. Wood (Pennsylvania State University, Pennsylvania, USA) (Hidalgo-Romano et al, 2014; Baba et al, 2006). Bacteria were cultured in LB medium until log phase. Subsequently, the log-phase bacteria were pelleted and re-suspended in fresh LB medium to an optical density of 1 at 600 nm (OD_600_ 1 = ∼10^9^ cfu/mL) for administration to GF mice.

### Metabolic chamber assessment

PhenoMaster (TSE Systems, Germany) was used for the metabolic chamber assessment. Briefly, the mice were transferred into the calorimetric chamber for immediate recording for 6 days, with the first 2.5 days serving as the habituation period, and the following 3 days for the analysis. Mice had *ad libitum* access to food and water and were under a 12 h light/dark cycle, a constant temperature of 22°C and humidity of 67%. Oxygen consumption, carbon dioxide production, food and water consumption, and body weight, were recorded for analysis.

### LC-MS targeted metabolomic analysis

All blood serum samples (10 WT and 10 MT) were prepared and analysed using a previously established method (Whileyet et al, 2019). In brief, 30 µL of serum was spiked with stable isotope labelled internal standards, prior to protein precipitation and removal which was performed using a two-step protocol consisting of addition of methanol followed bypass-through solid phase extraction. Samples were then taken to dryness under nitrogen and re-suspended in 10 mM ammonium formate with 0.5 % formic acid. 5 µL was then injected onto a Waters Acquity UHPLC system coupled to a Waters TQ-S tandem mass spectrometer (UHPLC-MS/MS). Chromatographic separation was performed on a reversed phase Waters Acquity HSS T3 C_18_ column (150mm x 2.1mm x 1.8µm) using an optimised 7 -minute solvent gradient.

The spectral data was pre-processed using TargetLynx (Waters, UK) to convert the LC-MS raw data into peak areas ratios with peak areas of internal standards. Calibration standards of a known concentration are used to produce a linear calibration plot. Quality control was performed using samples prepared with known concentrations of each metabolite and were used to assess accuracy and precision of the analysis throughout the data acquisition.

### NMR sample preparation, data acquisition and data analysis

NMR was performed according to an established protocol (Dona et al,2014). For plasma preparation, 100 μL of plasma was mixed with 50 μL of serum buffer (75 mM NaH_2_HPO_4_, pH=7.4, 100% D_2_O, 2 mM sodium azide and 0.08% TSP sodium salt), vortexed and 150 μL samples were transferred into a 3 mm NMR tube. NMR spectra were collected on a Bruker AVANCE III NMR spectrometer operating on 600.13 MHz with a BBI probe (Bruker Biospin, Germany) at 310 K. For each sample, a Carr-Purcell-Meiboom-Gill spectrum was obtained with the sequence ((RD-90°-(τ-180°-τ) n-acquisition); τ = 300 μs, n = 128). The relaxation delay (RD) was of 4 s, a 90° pulse was set at 7.6 μs (−11.03 db), and 32 free induction decays (FIDs) used 72K data-points and 20-ppm spectral width. For all spectral acquisition, the free induction decay NMR signals were multiplied by an exponential factor to give a line broadening of 0.3 Hz and were Fourier-transformed to obtain the usual frequency spectrum (TOPSPIN 3.5 software, Bruker Biospin, Rheinstetten, Germany). The spectra were automatically phased, baseline corrected and calibrated using the glucose signal at δ 5.23 ppm. ^1^H NMR spectral peaks were analysed using the student *t*-test for univariate analysis with *p*<0.05 being regarded as significant.

### Histological analysis by haematoxylin and eosin (H&E) staining

The intestine, adipose, and liver tissues were fixed in 4% formaldehyde, embedded in paraffin and sectioned for histological study. The intestines were Swiss-rolled before paraffin-embedding. Similar portions of the jejuna were used for analysis in all cases. Tissue sections were cut at 4 μm thickness with a microtome (Leica HistoCore MULTICUT) and stained with haematoxylin and eosin (H&E). Brightfield images of the sections were obtained using a Zeiss Axio Scan.Z1 slide scanner with 20x magnification, and subsequently analysed with a Zeiss ZEN Desktop microscope software.

### Immunohistochemistry

Paraffin-embedded intestine tissue sections were deparaffinized, rehydrated, and heat-treated with an antigen-retrieval buffer (sodium citrate buffer or Tris-EDTA-Tween 20 buffer). For KI67 labelling, sections were incubated with a rabbit polyclonal antibody (diluted 1:800, Abcam ab15580); For CHGA labelling, sections were incubated with another rabbit polyclonal antibody (diluted 1:800, Abcam ab45179). Sections were then incubated with ImmPRESS reagent (Vector, HRP-conjugated secondary antibodies and DAB substrate (Vector SK-4100), and counterstained with haematoxylin. Sections were imaged with a Zeiss Axio Scan.Z1 slide scanner with a 20x objective and analysed with a Zeiss ZEN software.

### Intestinal goblet cell quantification

Paraffin-embedded tissue sections were deparaffinized, rehydrated, and stained with periodic acid-Schiff (PAS) reagents. Haematoxylin were used for counterstaining of the nuclei. Sections were imaged with a Zeiss Axio Scan.Z1 slide scanner with a 20x objective and analysed with a Zeiss ZEN software. PAS-stained goblet cells were counted per villus.

### Liver oil red O staining

Liver tissues were fixed with 4% paraformaldehyde, equilibrated in 20% and 30% sucrose solution, and sectioned at 8 μm thickness with a Leica CM3050-S cryostat. The tissue sections were rinsed with 60% isopropanol, stained with freshly prepared oil red O (Sigma-Aldrich O0625-25G) solution, and lightly counter-stained with haematoxylin. Stained sections were imaged with a Zeiss Axio Scan.Z1 slide scanner with 20x objective and analysed with a Zeiss ZEN software.

### Quantitative real-time PCR

Total RNA was extracted from tissues with Qiagen RNeasy Mini Kit (Qiagen 74106) and used for complementary DNA synthesis with iScript Reverse Transcription Supermix (Bio-Rad 1708840), according to the manufacturers’ protocols. mRNA expression was quantified with Applied Biosystem StepOnePlus Real-Time PCR Systems, using Fast SYBR Green Master Mix (Applied Biosystems 4385612) or Roche Lightcycler 96, using the iTaq SYBR Green one-step kit (BioRad) and DNA oligo primers for targeted genes (primer sequences were tabulated in Table S2). At least 6 biological replicates were used for analysis with technical triplicates. Mouse *Hprt1* were used as the reference genes for expression normalization. DNA oligo primer sequences used are listed in Supplemental Figure 6.

### Western blotting

Tissues were homogenized in RIPA lysis buffer (Abcam ab156034, supplemented with phosphatase inhibitor cocktail, Roche 04906845001) for protein extraction. Bradford protein assay was used to determine protein concentration with standard curves established by BSA solution. Protein extract of fixed amount was subjected to electrophoresis and transferred to nitrocellulose membranes. Target proteins were detected by various primary antibodies (For phosphor-CREB rabbit mAb, Cell Signaling Technology #9198; For CREB rabbit mAb, Cell Signaling Technology #4820; For normalizing control β-Actin mouse mAb, Santa Cruz sc-47778), with HRP-conjugated secondary antibodies (Goat Anti-Rabbit Immunoglobulins/HRP, Dako P0448; Horse Anti-Mouse IgG, HRP-linked, Cell Signaling Technology #7076). BCIP/NBT substrate was used to detect. Results were quantified with a Bio-Rad ChemiDoc Imaging System and analysed by ImageJ software.

## Acknowledgments

We thank the following for funding support: Sunway University, Malaysia; National Neuroscience Institute, Singapore; Canadian Institute for Advanced Research, Canada (S.P.). This work is supported by the UK Dementia Research Institute [award number UKDRI-5002] which receives its funding from UK DRI Ltd, funded by the UK Medical Research Council, Alzheimer’s Society and Alzheimer’s Research UK (A.T.B). The authors would like to acknowledge the financial support from National Research Foundation and Ministry of Education Singapore under its Research Centre of Excellence Program and NIMBLES grant (S.A.R). We thank Dr Hae Ung Lee, Ms Alicia Kang, Dr Zoe Bichler and Dr Jintao Wang for their kind assistance.

## Author contributions

P.X.Y and S.P. contributed to the conception and design of the study; P.X.Y., A.J., G.W.Z., K.A.M., L.E.F., Y.W., A.T.B., E.H., J.N., and L.W. contributed to the acquisition and analyses of data; P.X.Y., A.J., and S.P., contributed to drafting the manuscript and preparing the figures; S.K., S.A.R., and S.P., reviewed the project.

## Summary

### PROBLEM

<u>G</u>ut microbes and their metabolites are known to regulate host organ growth and function. One such microbe regulating pathway is the metabolites generated from tryptophan metabolism – planar indoles. Here we assessed the impact of indole on host organ growth and function.

### RESULTS

We used germ-free male mice mono-colonized with indole-producing wildtype *Escherichia coli* or tryptophanase-encoding *tnaA* knockout mutant indole-non-producing *E. coli*. Compared to control mice, the mutant *E. coli* recipient mice displayed a severe wasting phenotype including multiorgan decline (muscle liver, adipose tissue), growth retardation, catabolism and energy deficiency. In addition, there were multiple deficiencies in the alimentary tract involving digestion problems and energy harvest, reduced bowel movement due to reduced serotonin levels and impairment in intestinal neuronal function. Collectively, these are consistent with accelerated organ decline, thus mimicking an accelerated ageing phenotype.

### IMPACT

Our results demonstrate that microbe-derived indoles are necessary to maintain adult metabolic homeostasis across multiple organs and suggest that reduced levels of indoles could hasten biological ageing and the risk for unwanted organ decline among human beings.

## SUPPLEMENTAL INFORMATION

**Supplemental Table 1.**
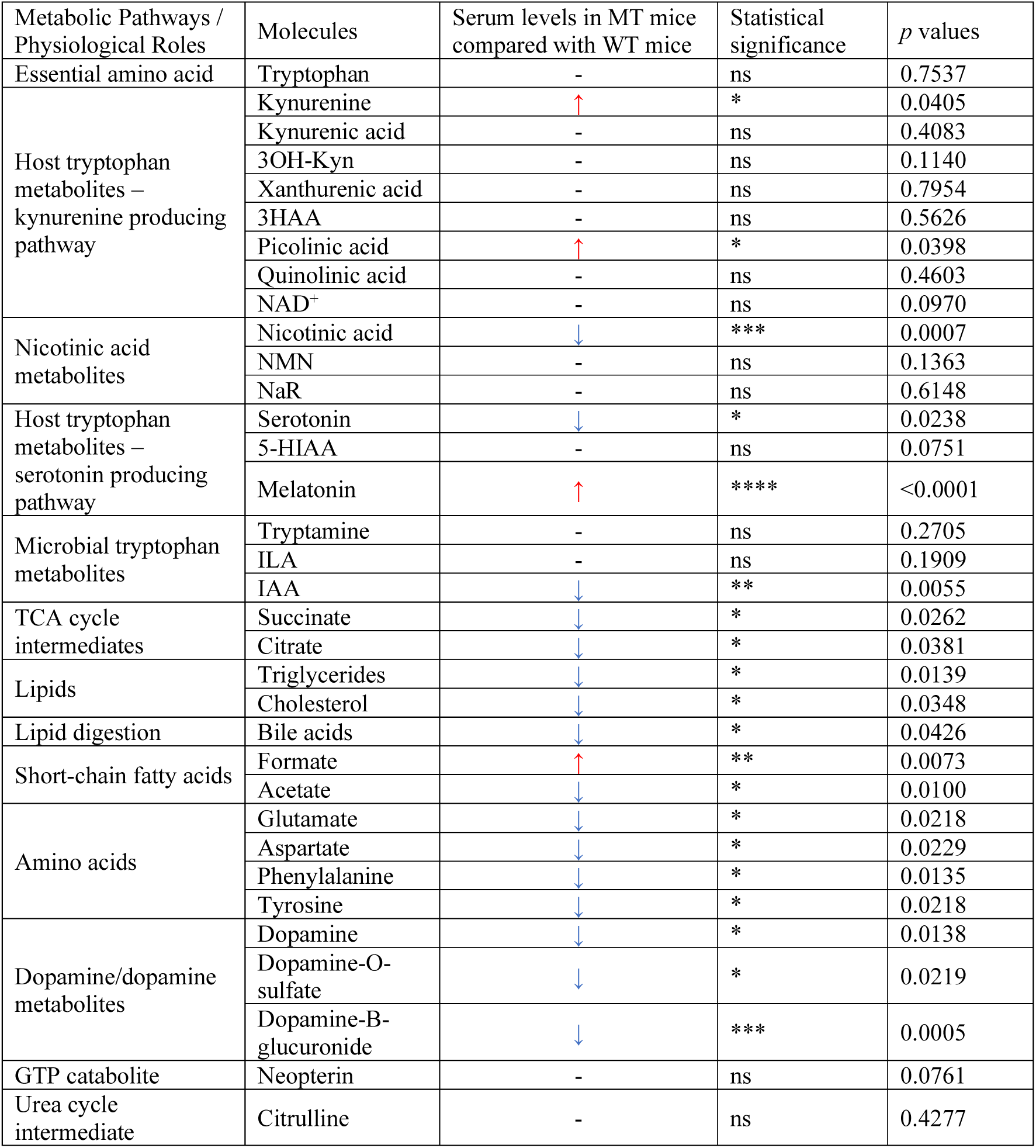
Serum levels of selected metabolites by metabolomic studies. n = 8-10 per group. *p* values were calculated with the student’s *t* test. Statistical significance is presented as \**p* < 0.05, \*\**p* < 0.01, \*\*\**p* < 0.001, and \*\*\*\**p* < 0.0001 between indicated groups.

**Supplemental Table 2.**
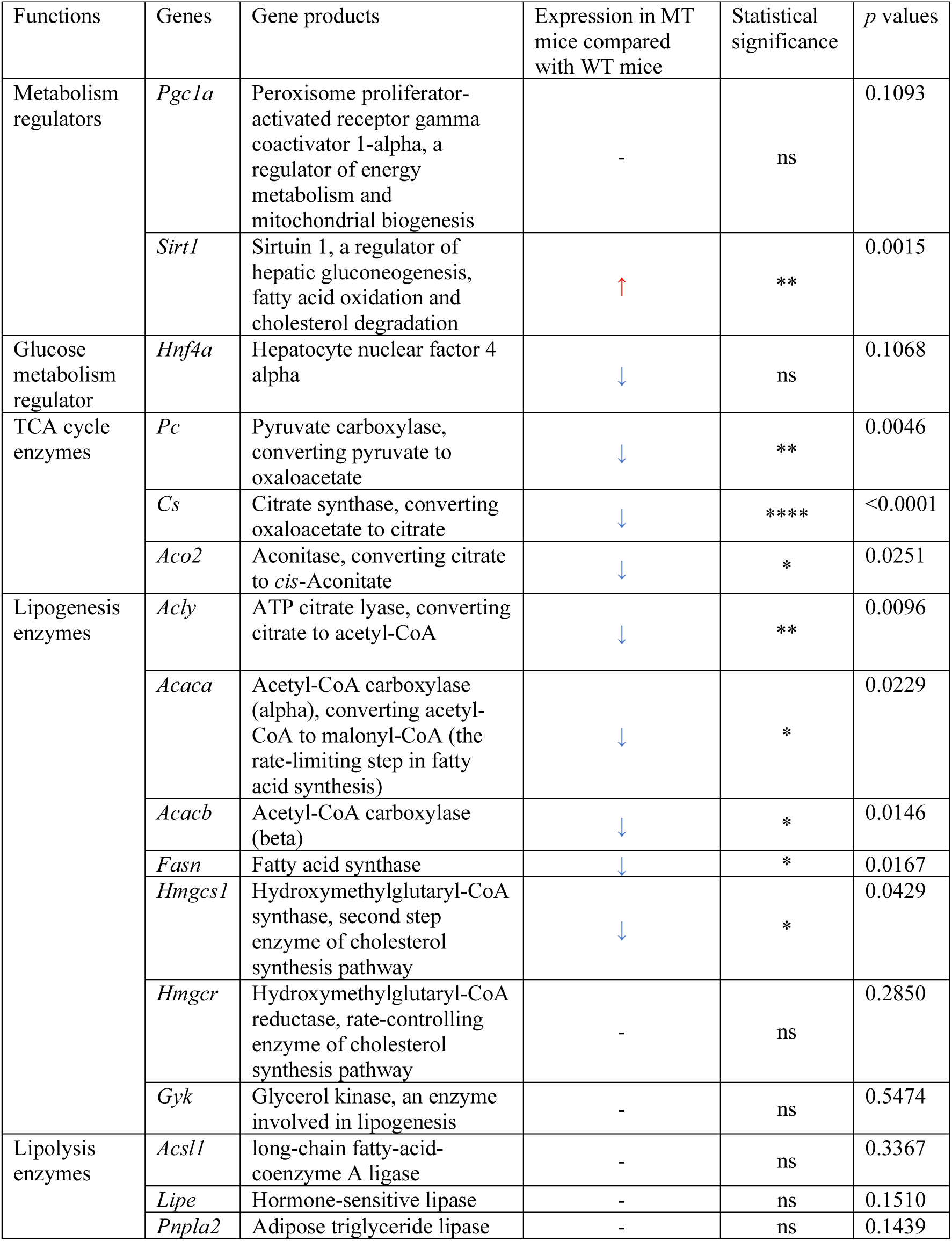
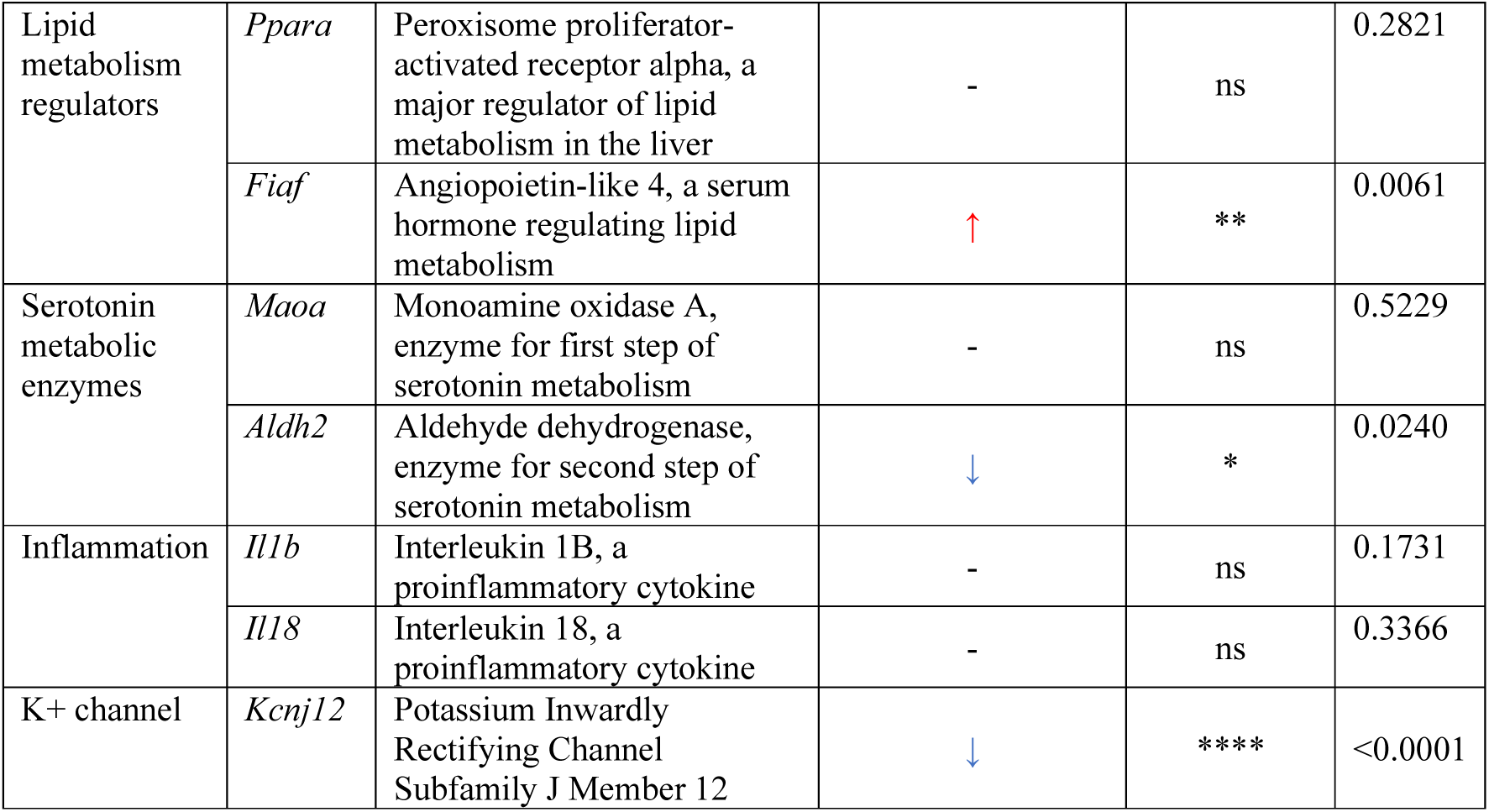
mRNA expression levels of selected genes in liver samples from WT and MT mice. n=7-8 per group. *p* values were calculated with the student’s *t* test. Statistical significance is presented as \**p* < 0.05, \*\**p* < 0.01, \*\*\**p* < 0.001, and \*\*\*\**p* < 0.0001 between indicated groups. ns denotes no statistical significance with *p* ≥ 0.05.

**Supplemental Table 3.**
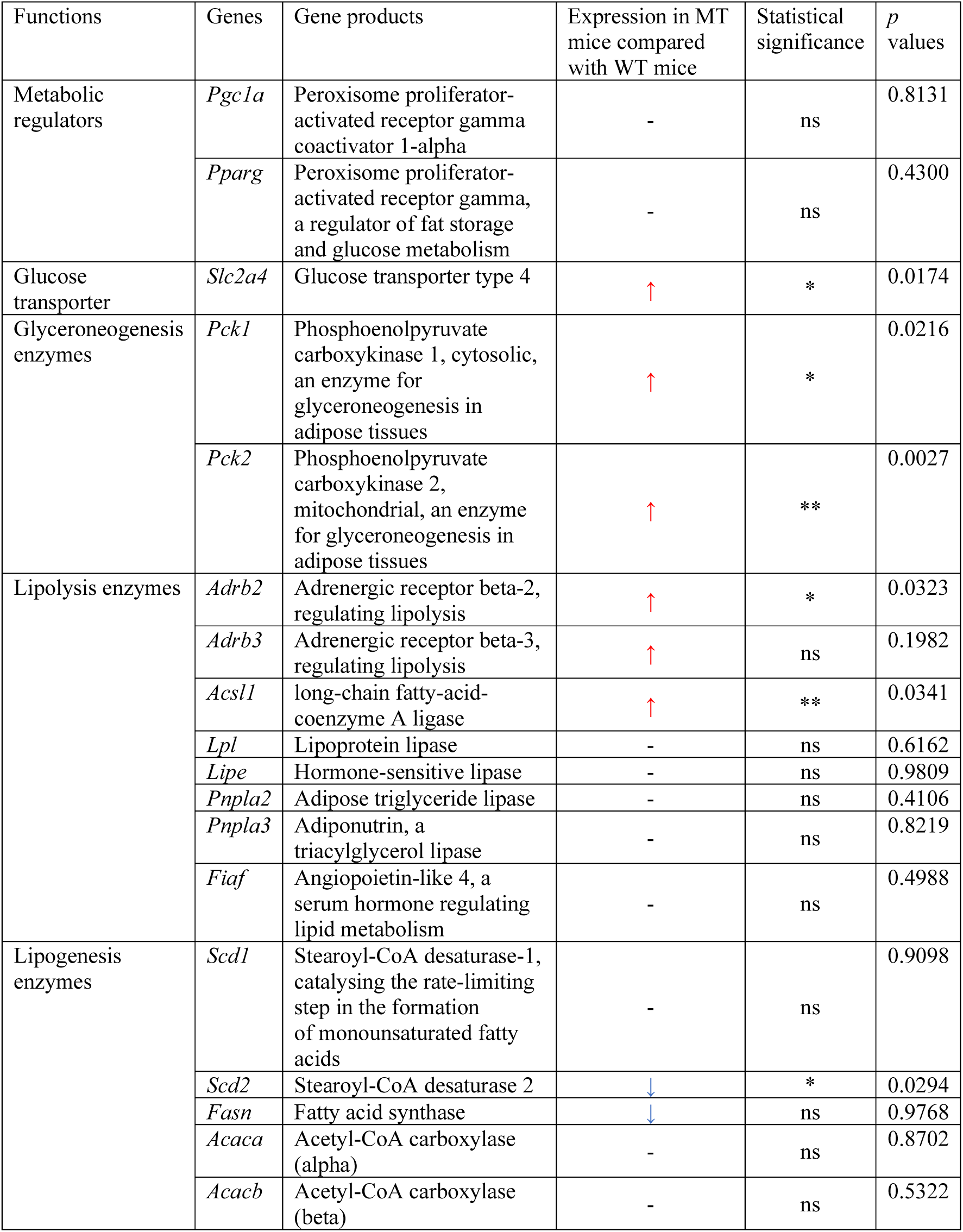
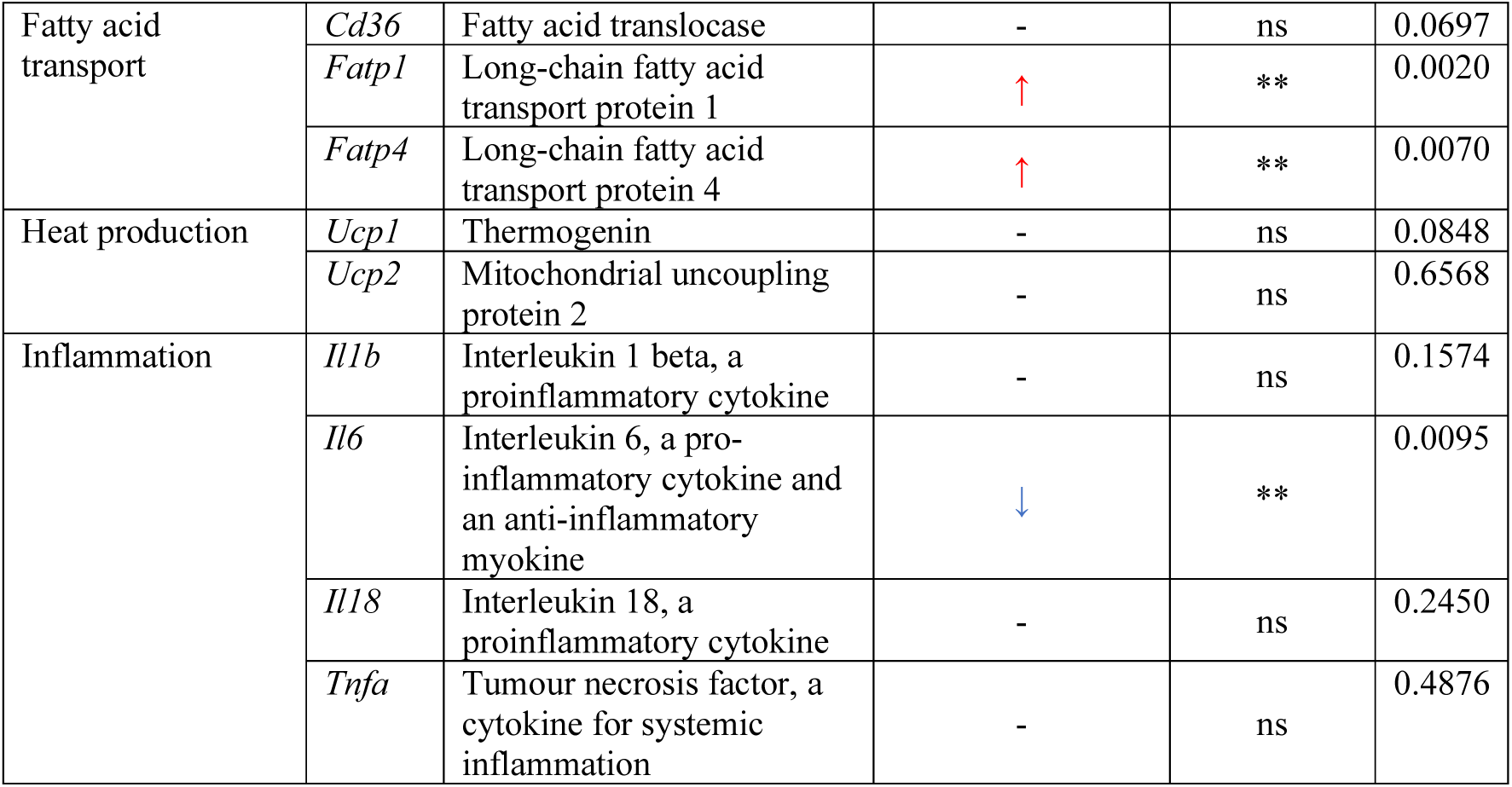
mRNA expression levels of selected genes in epididymal white adipose tissue (WAT) samples from WT and MT mice. n=6-8 per group. *p* values were calculated with the student’s *t* test. Statistical significance is presented as \**p* < 0.05, \*\**p* < 0.01, \*\*\**p* < 0.001, and \*\*\*\**p* < 0.0001 between indicated groups. ns denotes no statistical significance with *p* ≥ 0.05.

**Supplemental Table 4.** mRNA expression levels of selected genes in hindlimb muscles gastrocnemius (GS) samples from WT and MT mice. n=7-8 per group. *p* values were calculated with the student’s *t* test. Statistical significance is presented as \**p* < 0.05, \*\**p* < 0.01, \*\*\**p* < 0.001, and \*\*\*\**p* < 0.0001 between indicated groups. ns denotes no statistical significance with *p* ≥ 0.05.

| Functions | Genes | Gene products | Expression in MT mice compared with WT mice | Statistical significance | *p* values |
| --- | --- | --- | --- | --- | --- |
| Metabolism regulators | *Pgc1a* | Peroxisome proliferator-activated receptor gamma coactivator 1-alpha, a regulator of energy metabolism and mitochondrial biogenesis | - | ns | 0.9843 |
|  | *Pparg* | Peroxisome proliferator-activated receptor gamma, a regulator of fat storage and glucose metabolism | - | ns | 0.3779 |
| Glucose metabolism and mitochondrial functions | *Glut4* | Glucose transporter | ↓ | * | 0.0285 |
|  | *Sdh* | succinate dehydrogenase, a TCA cycle and respiratory chain enzyme | ↓ | ns | 0.1879 |
|  | *Tfam* | Mitochondrial transcription factor A | ↓ | * | 0.0456 |
|  | *Pdk4* | Pyruvate dehydrogenase lipoamide kinase isozyme 4, mitochondrial, a regulator of glucose metabolism | - | ns | 0.6495 |
| Lipid metabolism | *Ppara* | Peroxisome proliferator-activated receptor alpha, a major regulator of lipid metabolism in the liver | - | ns | 0.6454 |
|  | *Fiaf* | Angiopoietin-like 4, a serum hormone regulating lipid metabolism | - | ns | 0.9469 |
|  | *Srebp2* | Sterol regulatory element-binding protein 2, a transcription factor regulating cholesterol homeostasis | ↓ | * | 0.0138 |
| Muscle components | *Myhc-1* | Myosin heavy chain 1 | - | ns | 0.5249 |
|  | *Myhc-2a* | Myosin heavy chain 2a | - | ns | 0.5336 |
|  | *Myhc-2b* | Myosin heavy chain 2b | - | ns | 0.4575 |
|  | *Myhc-2d* | Myosin heavy chain 2d | ↓ | * | 0.0182 |
|  | *Rapsyn* | Receptor-associated protein of the synapse (neuromuscular junction-associated gene) | ↓ | ns | 0.2569 |
| Muscle atrophy | *Murf-1* | Muscle RING-finger protein-1, a muscle atrophy E3 ubiquitin ligase | ↓ | * | 0.0215 |
|  | *Atrogin-1* | A muscle atrophy E3 ubiquitin ligase | ↓ | ** | 0.0055 |
| Myogenesis | *Myod* | Myoblast determination protein 1, regulating muscle differentiation | ↓ | n.s. | 0.1975 |
| Inflammation | *Il6* | Interleukin 6, a pro-inflammatory cytokine and an anti-inflammatory myokine | - | ns | 0.4321 |
| K<sup>+</sup> channel | *Kcnj12* | Potassium Inwardly Rectifying Channel Subfamily J Member 12 | - | ns | 0.3598 |

**Supplemental Table 5.** mRNA expression levels of selected genes in small intestine (SI) samples from WT and MT mice. n=7-8 per group. *p* values were calculated with the student’s *t* test. Statistical significance is presented as \**p* < 0.05, \*\**p* < 0.01, \*\*\**p* < 0.001, and \*\*\*\**p* < 0.0001 between indicated groups. ns denotes no statistical significance with *p* ≥ 0.05.

| Functions | Genes | Gene products | Expression in MT mice compared with WT mice | Statistical significance | *p* values |
| --- | --- | --- | --- | --- | --- |
| Gut barrier integrity | *Occludin* | Occludin, an epithelial cell tight junction protein | - | ns | 0.1124 |
|  | *Zo-1* | Zonula occludens-1, an epithelial cell tight junction protein | - | ns | 0.6254 |
|  | *Claudin-1* | Claudin-1, an epithelial cell tight junction protein | ↑ | ns | 0.0636 |
| Inflammation | *Tnfa* | Tumour necrosis factor, a cytokine for systemic inflammation | ↑ | ns | 0.2304 |
|  | *Il1b* | Interleukin 1 beta, a proinflammatory cytokine | ↑ | * | 0.0443 |
|  | *Il18* | Interleukin 18, a proinflammatory cytokine | - | ns | 0.1732 |
|  | *Cd68* | Protein expressed by immune cells | - | ns | 0.5714 |
| Serotonin synthesis, transport, and metabolism | *Tph1* | Tryptophan hydroxylase 1, first-step enzyme in serotonin synthesis | - | ns | 0.5121 |
|  | *Ddc* | Aromatic amino acid decarboxylase, second-step enzyme in serotonin synthesis | - | ns | 0.5296 |
|  | *Slc6a4* | Serotonin transporter | - | ns | 0.0962 |
|  | *Maoa* | Monoamine oxidase A, enzyme for first step of serotonin metabolism | - | ns | 0.9137 |
|  | *Aldh2* | Aldehyde dehydrogenase, enzyme for second step of serotonin metabolism | - | ns | 0.0934 |
| Active intestinal stem cell markers | *Lgr5* | Leucine-rich G-protein-coupled receptor 5 | ↓ | *** | 0.0007 |
|  | *Olfm4* | Olfactomedin 4 | - | ns | 0.2988 |
| Goblet cell marker | *Muc2* | Mucin 2 | - | ns | 0.2306 |
| Wnt signalling pathway (regulating stem cell self-renewal) | *Ctnnb1* | β-catenin, a protein involved in Wnt signalling, a pathway important for intestinal stem cell renewal | - | ns | 0.3537 |
|  | *Lrp6* | Low-density lipoprotein receptor-related protein 6, a receptor in Wnt signalling | - | ns | 0.7782 |
|  | *Fzd7* | Frizzled-7, a receptor in Wnt signalling | - | ns | 0.2633 |
| Notch signalling (modulating stem cell differentiation) | *Hes1* | A transcription factor interacts with Notch signalling, modulating secretory lineage differentiation in intestinal epithelium | - | ns | 0.2309 |
| Cell proliferation | *Mcc* | Colorectal mutant cancer protein, a negative regulator of cell cycle progression | ↓ | * | 0.0171 |
|  | *Pcna* | Proliferating cell nuclear antigen | - | ns | 0.0587 |
|  | *cMyc* | A transcription factor regulating many pro-proliferation genes | - | ns | 0.6450 |
|  | *Ccnd1* | Cyclin D1, a protein required for progression through G1 phase of cell cycle | - | ns | 0.4143 |
|  | *Cdkn1a* | Cyclin-dependent kinase (CDK) inhibitor 1, p21Cip1, inhibitor of cyclins/CDKs, including PCNA | - | ns | 0.7205 |
| Glucose metabolism | *Sirt1* | Sirtuin 1, a regulator of hepatic gluconeogenesis, fatty acid oxidation and cholesterol degradation | - | ns | 0.7542 |
|  | *Glut2* | Glucose transporter 2 | - | ns | 0.5875 |
|  | *Abcg5* | Proteins for regulation of sterol absorption and excretion | - | ns | 0.7158 |
|  | *Abcg8* |  | - | ns | 0.1150 |
|  | *Idh2* | Isocitrate dehydrogenase [NADP], mitochondrial, a TCA cycle enzyme | ↓ | ** | 0.0046 |
| Lipid metabolism | *Npc1l1* | A protein involved in cholesterol absorption | ↓ | * | 0.0175 |
|  | *Scarb1* | Scavenger receptor class B type 1, a receptor involved in cholesterol transport | - | ns | 0.5271 |
|  | *Fatp4* | Long-chain fatty acid transport protein 4 | - | ns | 0.9533 |
| Neural signalling | *Npy* | Neuropeptide Y, modulating diverse neurological processes including food intake and fat storage | - | ns | 0.2131 |
| K<sup>+</sup> channel | *Kcnj12* | Potassium Inwardly Rectifying Channel Subfamily J Member 12 | ↓ | ** | 0.004 |

**Supplemental Table 6.**
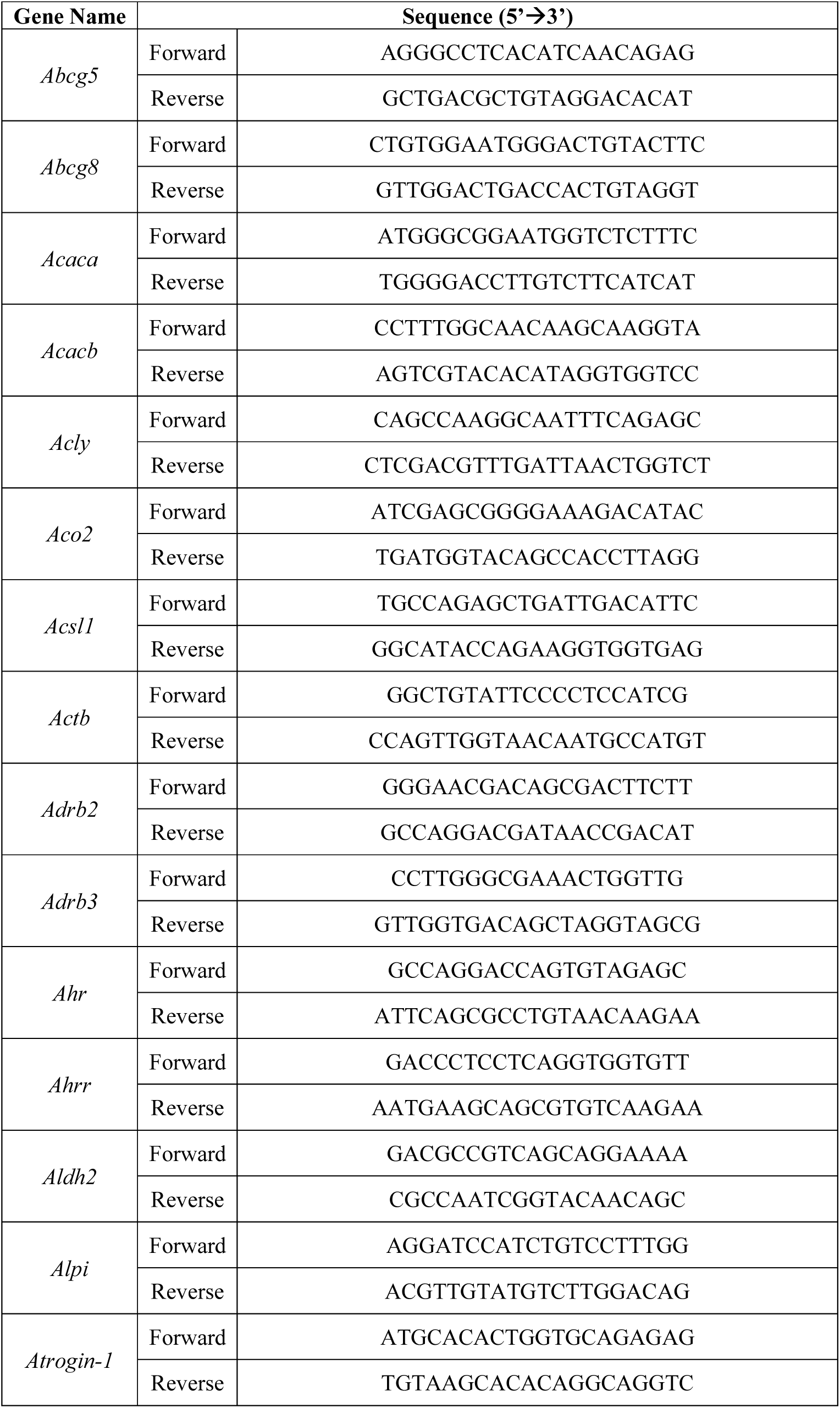
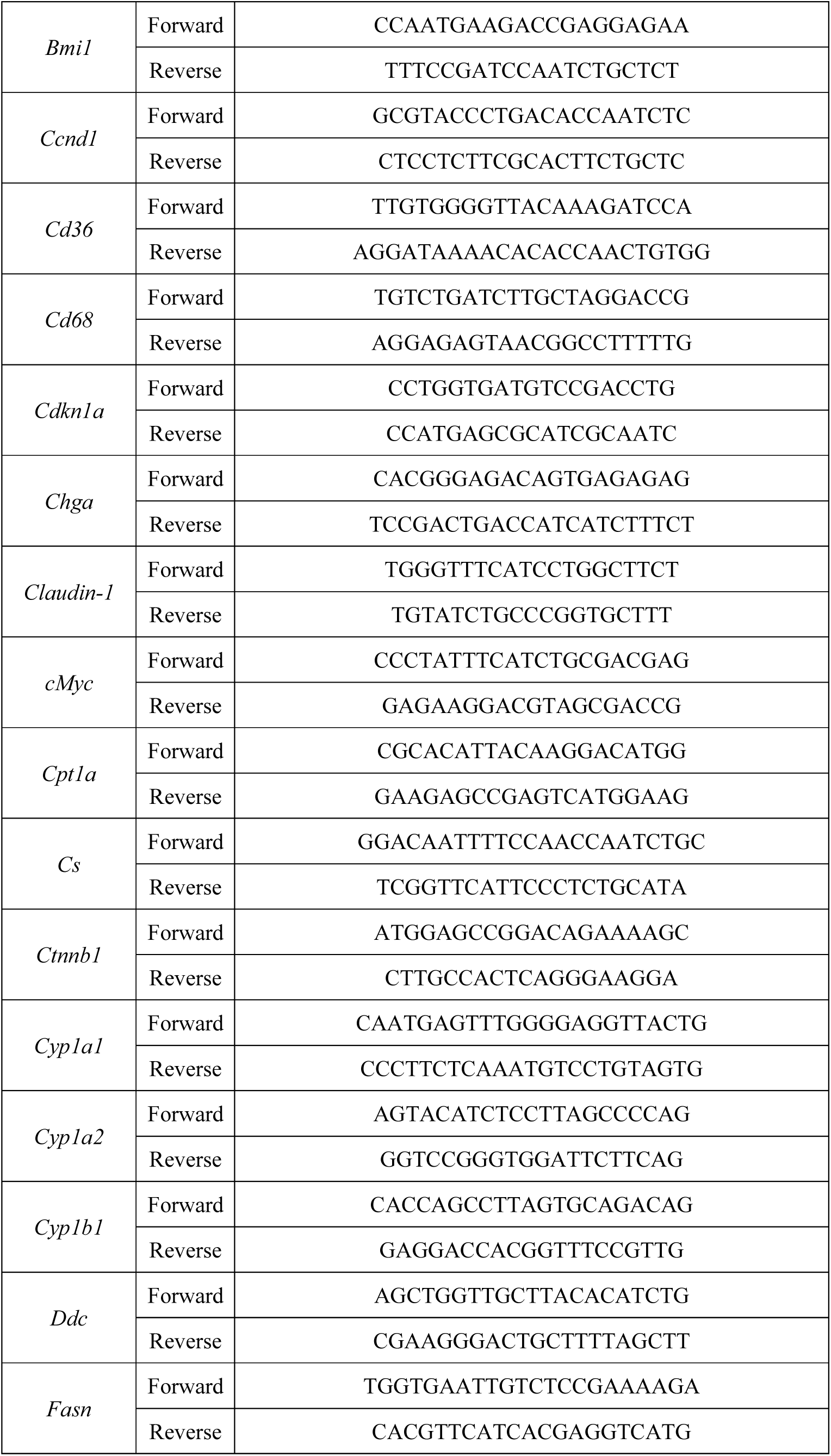
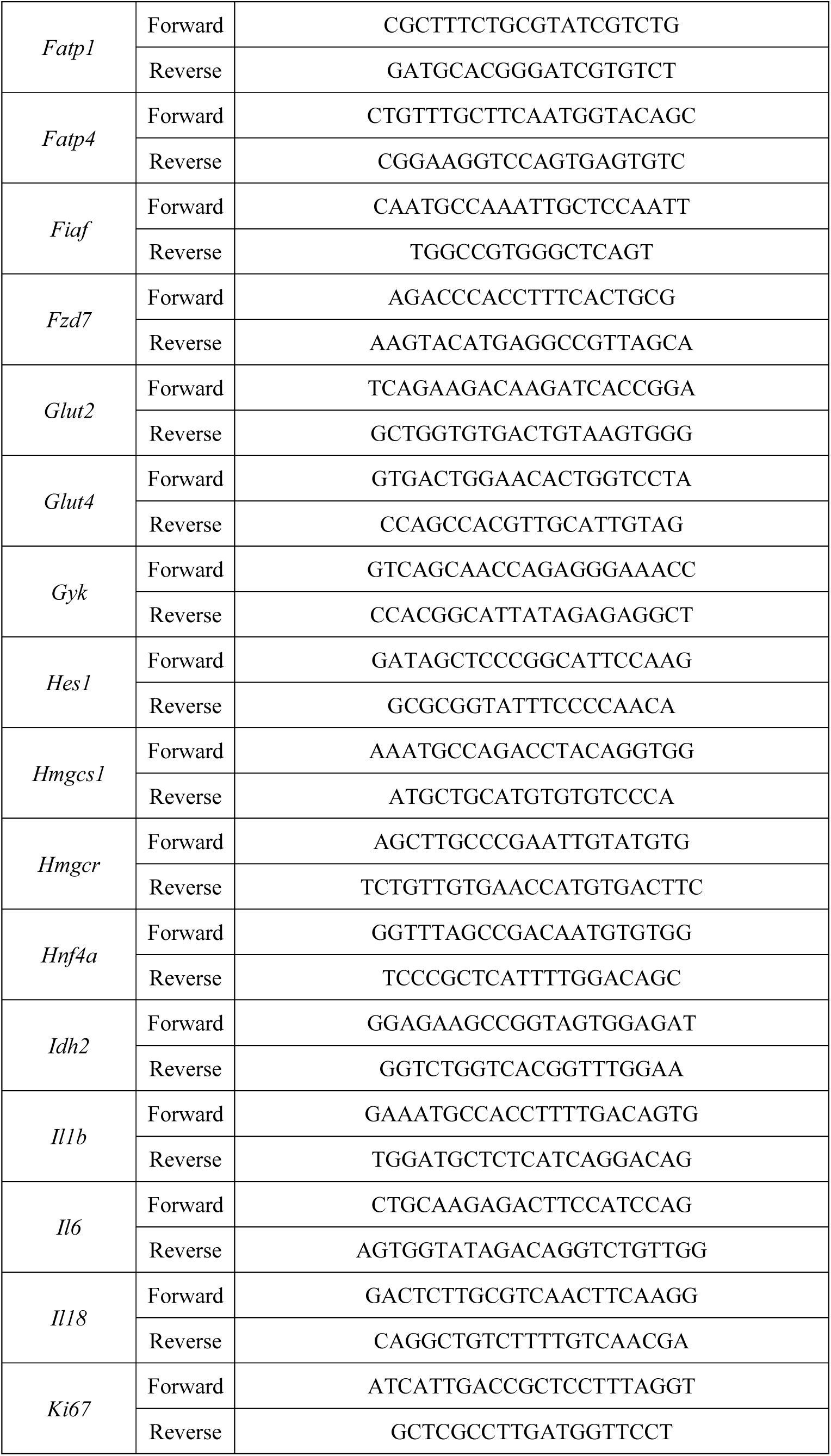
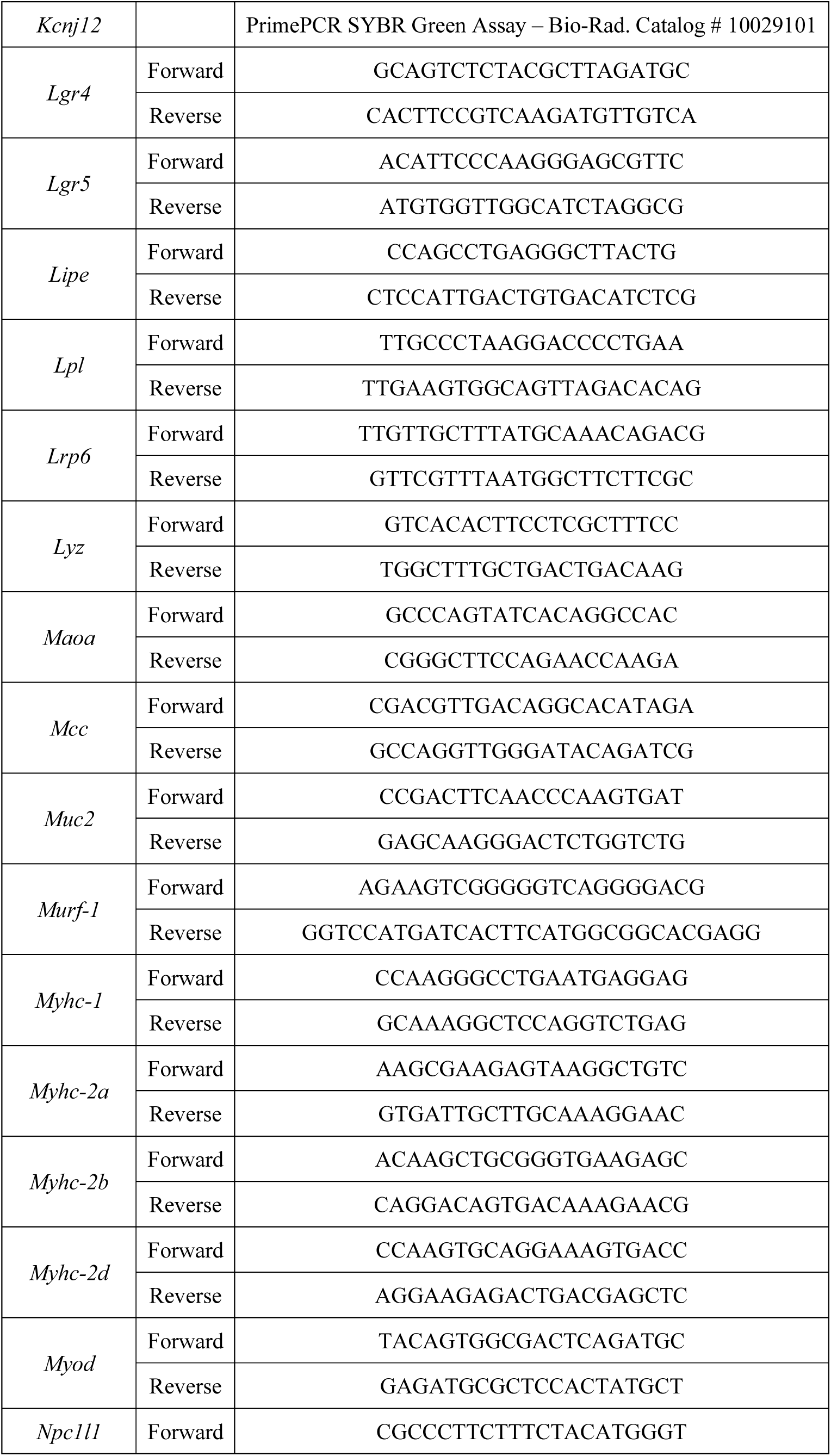
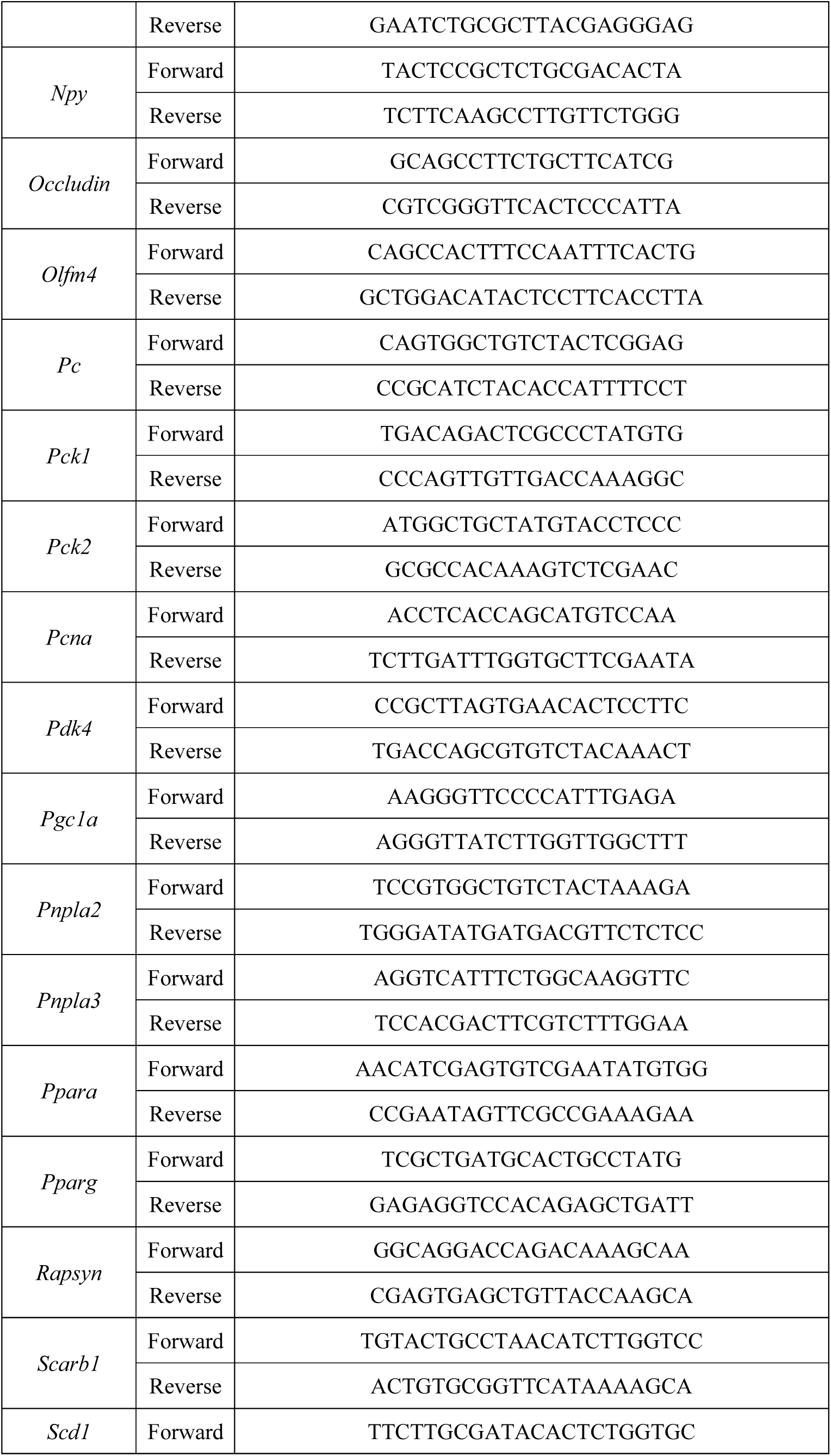
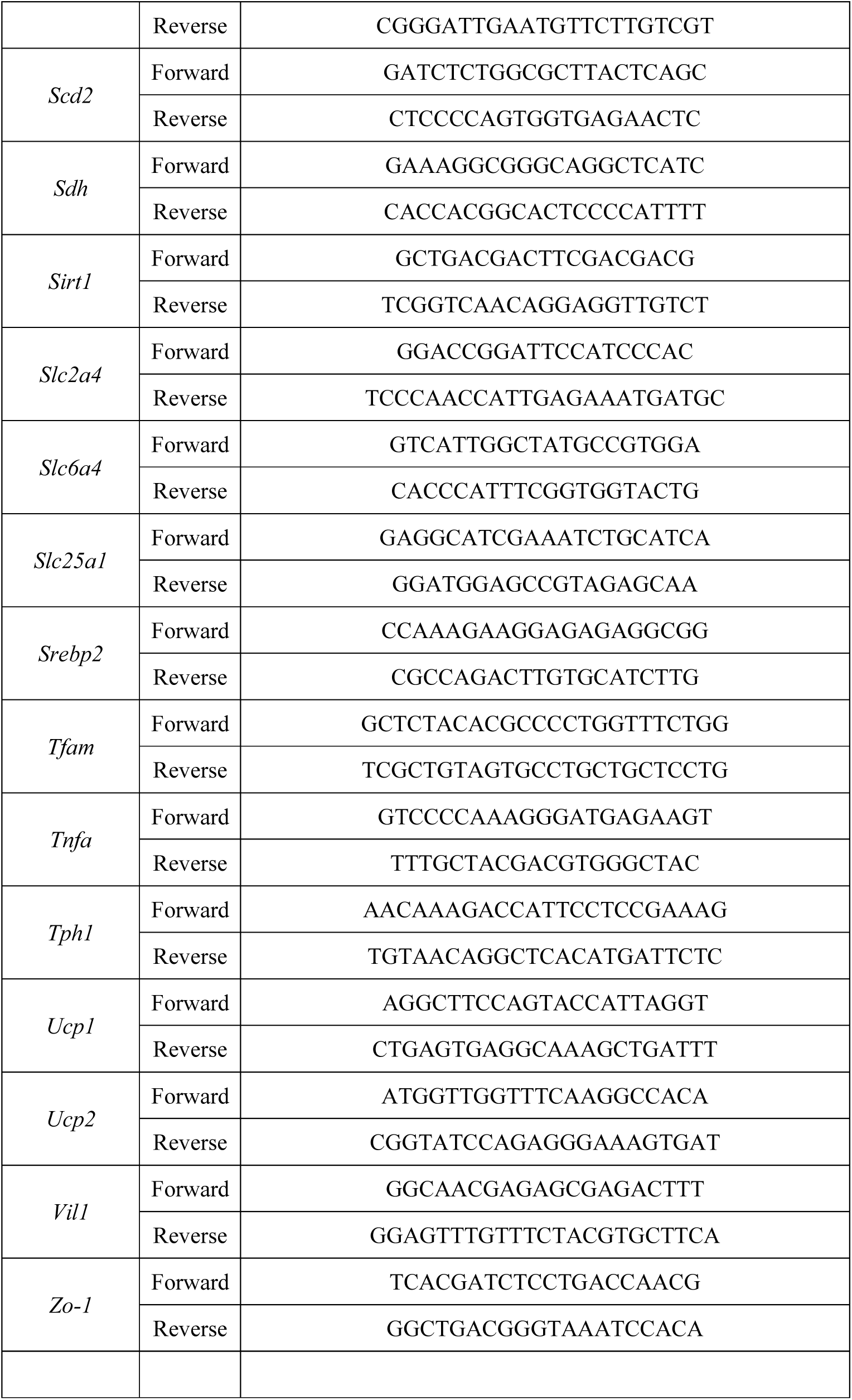
DNA Oligo Primer Sequences Used in qPCR.

**Supplemental Figure 1.**
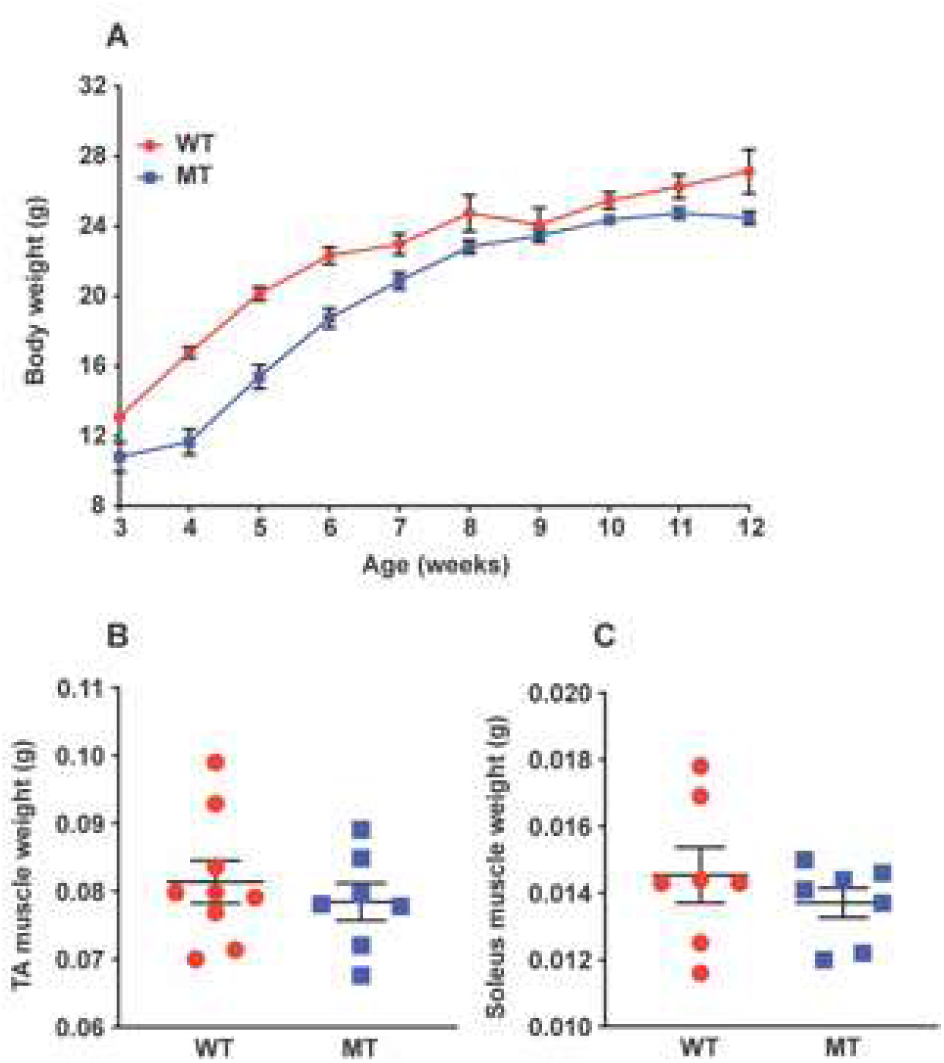
**A** Body weights of WT and MT mice from weaning (3 weeks old) to 12 weeks old. (n = 6-21 at each time point per group) **B**, **C** Weights of WT and MT mouse hindlimb muscles tibialis anterior (TA, **B**) and soleus (**C**) (n = 7-10 per group). Data graphs show means ± SEM error bars.

**Supplemental Figure 2.**
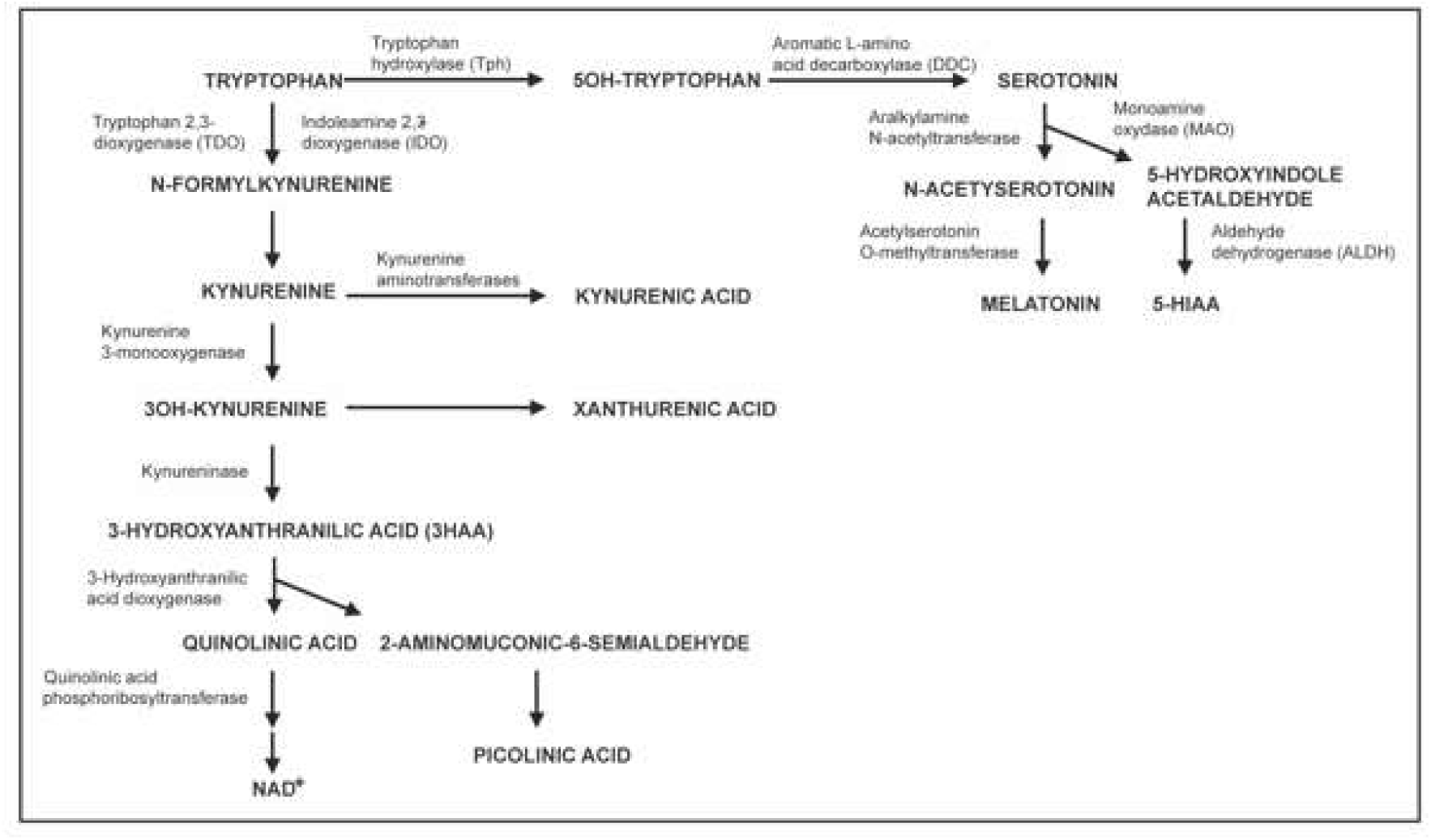
Illustration of tryptophan metabolisms in the mammalian host. Tryptophan can be metabolized in the host via the kynurenine pathway, through which kynurenine and NAD^+^, along with other metabolites, are produced. Alternatively, tryptophan can also be metabolized in the intestines by the enterochromaffin cells, or in the brain, through the serotonin pathway, leading to the production of serotonin. Serotonin can be further metabolized to 5-hydroxyindoleacetic acid (5-HIAA) or be utilized as a substrate for production of melatonin, a hormone regulating the sleep-wake cycle. NAD^+^, nicotinamide adenine dinucleotide (oxidized). Figure is adapted from Schwarcz & Stone, 2017. Copyright © 2016 Elsevier Ltd. Adapted with permission.

**Supplemental Figure 3.**
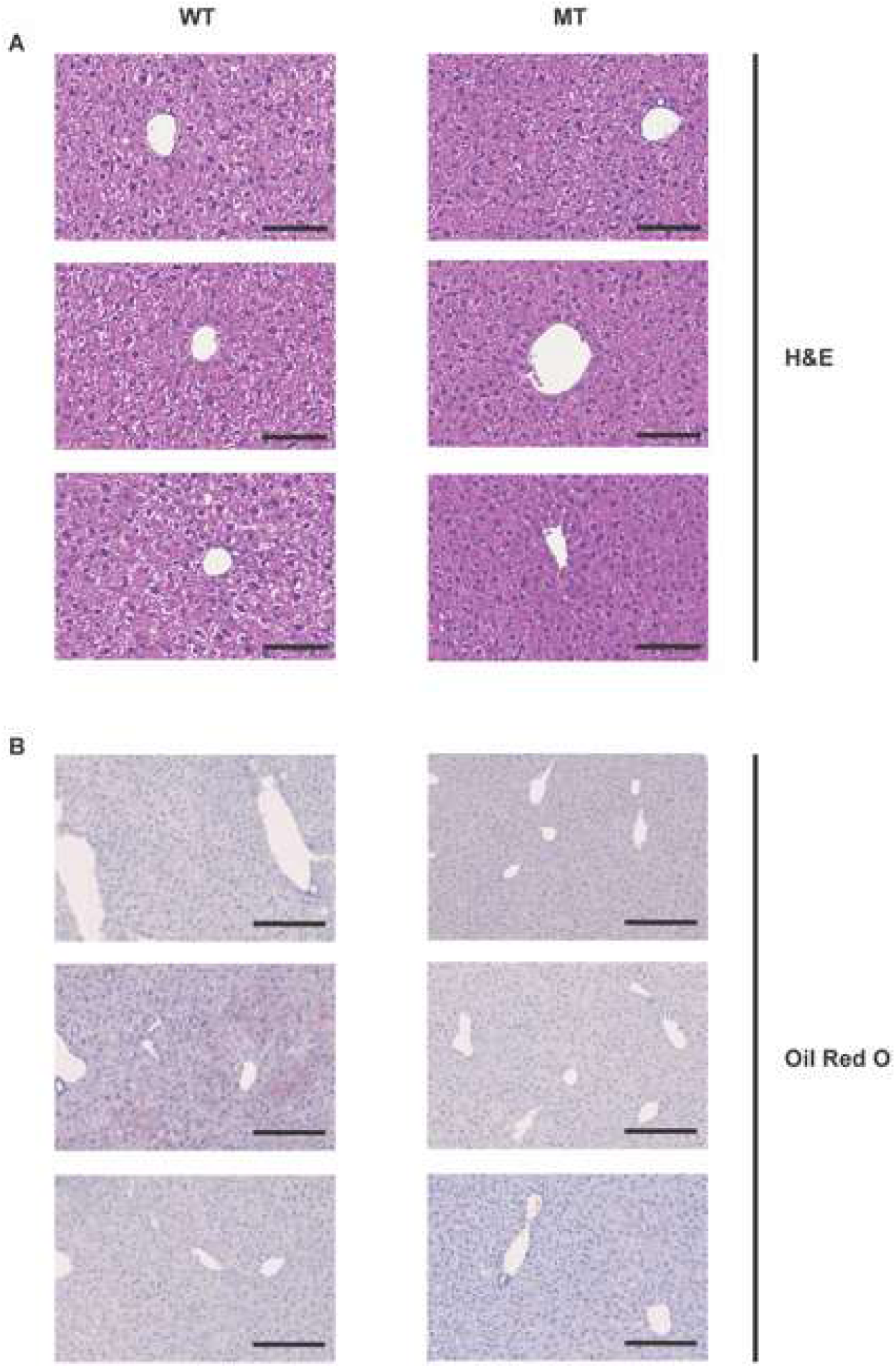
Representative images of liver histological analyses by (**A**) haematoxylin and eosin (H&E) and (**B**) oil red O staining. For **A**, scale bars = 100 μm; For **B**, scale bars = 200 μm.

**Supplemental Figure 4.**
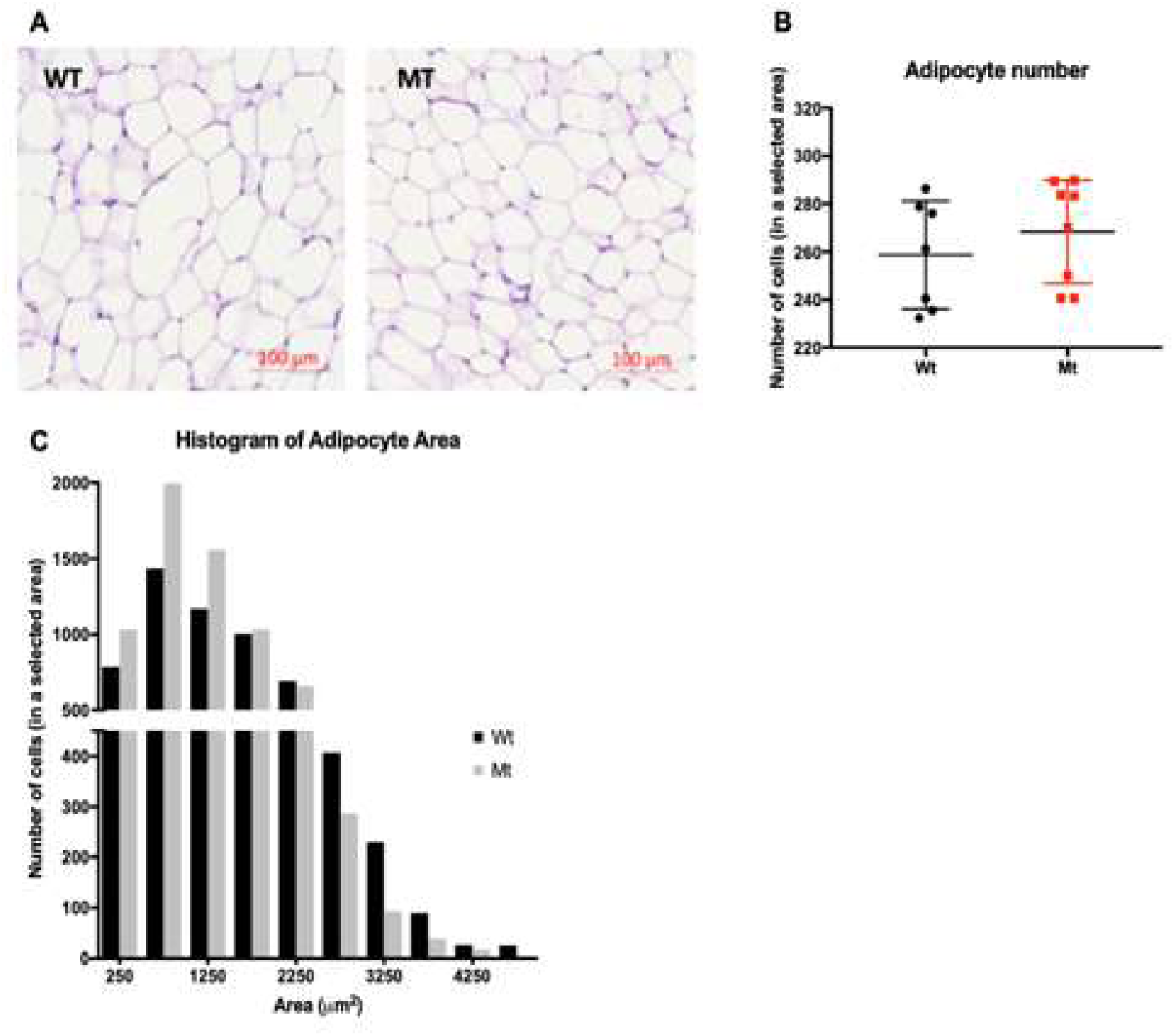
Epididymal white adipose tissue (WAT) morphology. **A** Representative images of WAT histological analyses by haematoxylin and eosin (H&E). Scale bars = 100 μm. **B** Adipocyte number in a selected area. **C** Histogram of adipocyte areas.

**Supplemental Figure 5.**
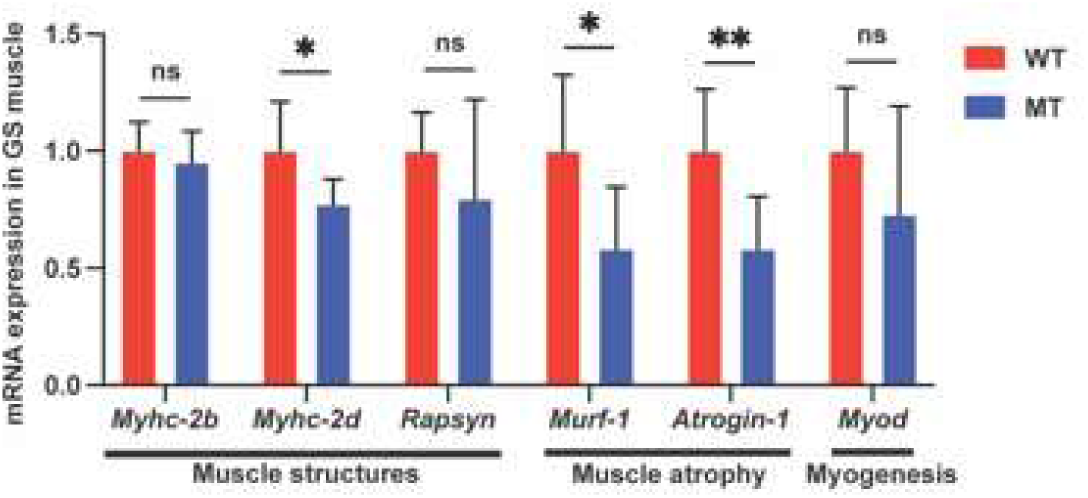
Indole-non-producing *E. coli*-colonized mice showed reduced expression levels of muscle atrophy-related and myogenesis-related genes in hindlimb muscles. mRNA expression of expression of myosin heavy-chain II proteins *Myhc-2b* and *Myhc-2d*, neuromuscular junction-associated gene *Rapsyn* (43 kDa receptor-associated protein of the synapse), muscle atrophy E3 ubiquitin ligases *Murf-1* (E3 ubiquitin-protein ligase TRIM63) and *Atrogin-1* (F-box only protein 32), and myogenesis-related gene *Myod* (myoblast determination protein 1) in GS muscles of WT and MT mice. (n=7-8 per group). mRNA levels were normalized to the control gene *Hprt1*. Data graphs show means ± S.D. *p* values were calculated with the student’s *t* test. Statistical significance is presented as \**p* < 0.05 and \*\**p* < 0.01 between indicated groups. ns denotes not significant with *p* ≥ 0.05.

**Supplemental Figure 6.**
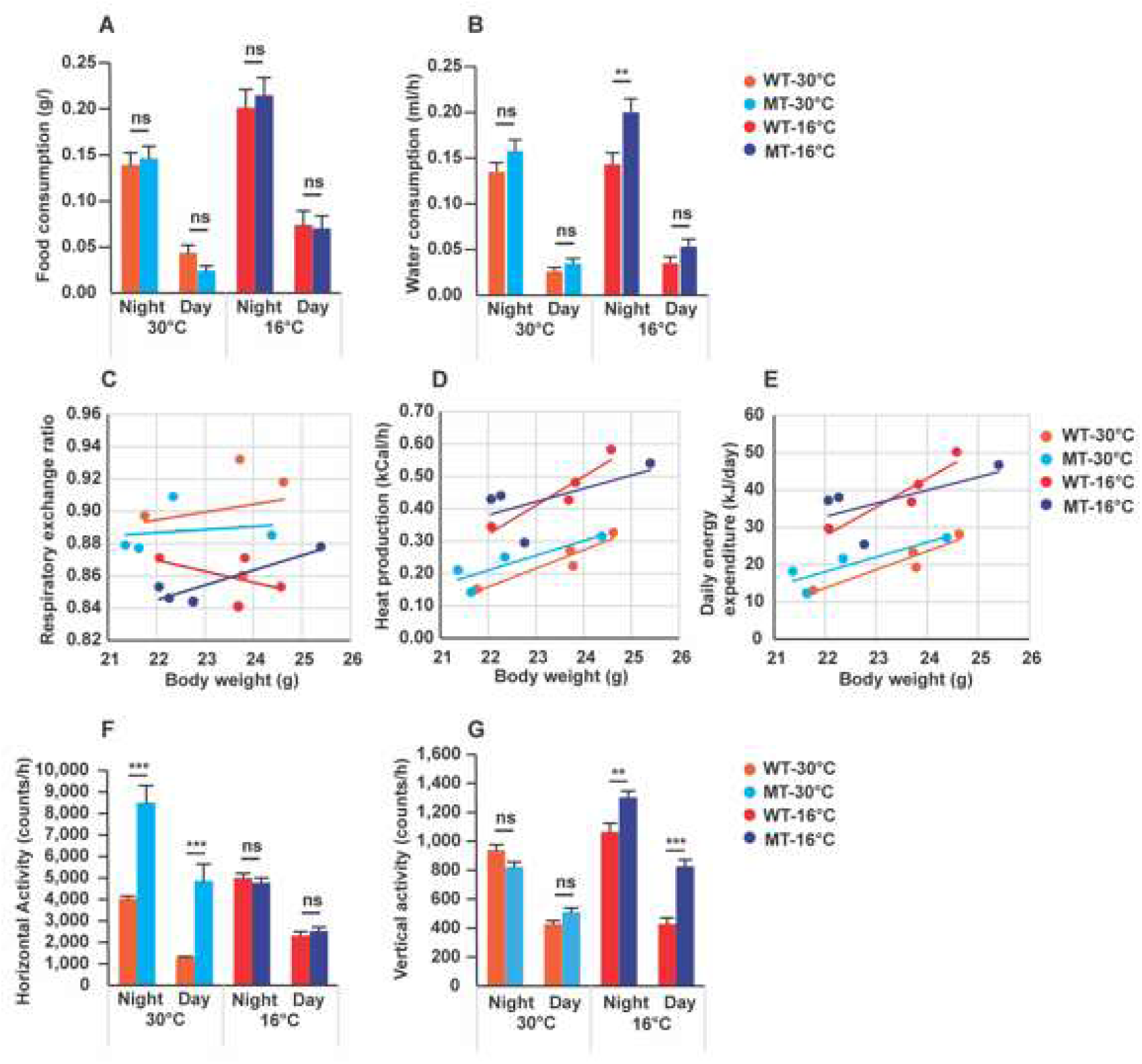
Calorimetric cage analyses at 30°C and 16°C for WT and MT mice. **A, B** Food and water consumption at night- and daytime of WT and MT mice at 30°C and 16°C. **C** to **E** Linear regression curves for correlation of body weight on the respiratory exchange rates, heat production, and daily energy expenditure. Data are the means of results over 3 experimental days per animal. Each dot represents an animal. **F, G** Horizontal and vertical activities of WT and MT mice. Experiments were at a constant temperature of 30°C or 16°C with humidity of 67%. A total of 3 days in the calorimetric cages were used for the data analysis. n=4 per group. For **A**, **B**,**F**,**G**, data graphs show means ± SEM error bars. *p* values were calculated with ANOVA (Welch’s robust *t* test of equality of means). Statistical significance is presented as \**p* < 0.05, \*\**p* < 0.01, \*\*\**p* < 0.001, and \*\*\*\**p* < 0.0001 between indicated groups. ns denotes not significant with *p* ≥ 0.05. For **C**, **D**,**E**, data were analysed by ANCOVA GLM statistical analysis comparing the 2 groups with the body weight and day/night variation as covariates.

**Supplemental Figure 7.**
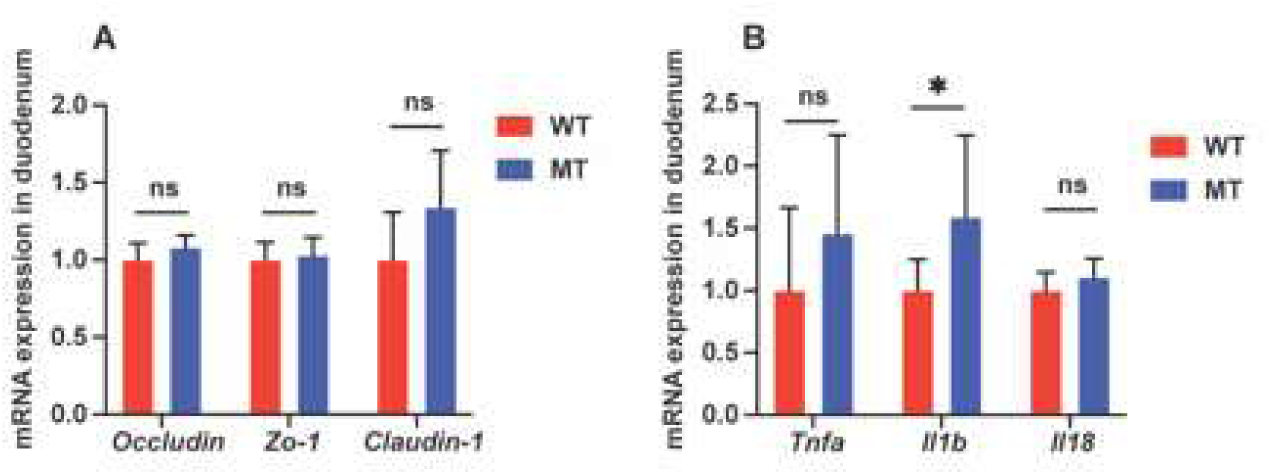
qPCR analyses on (**A**) intestinal tight junction proteins, *Occludin*, *Zo-1* (zonula occludens-1), and *Claudin-1* (n=8 per group); and (**B**) pro-inflammatory cytokines *Tnfa* (tumour necrosis factor alpha), *Il1b* (interleukin 1 beta), *Il18* (interleukin 18), in the duodena of WT and MT mice (n=7-8 per group). mRNA levels were normalized to the control gene *Hprt1*. Data graphs show means ± S.D. *p* values were calculated with the student’s *t* test. Statistical significance is presented as \**p* < 0.05 between indicated groups. ns denotes not significant with *p* ≥ 0.05.

**Supplemental Figure 8.**
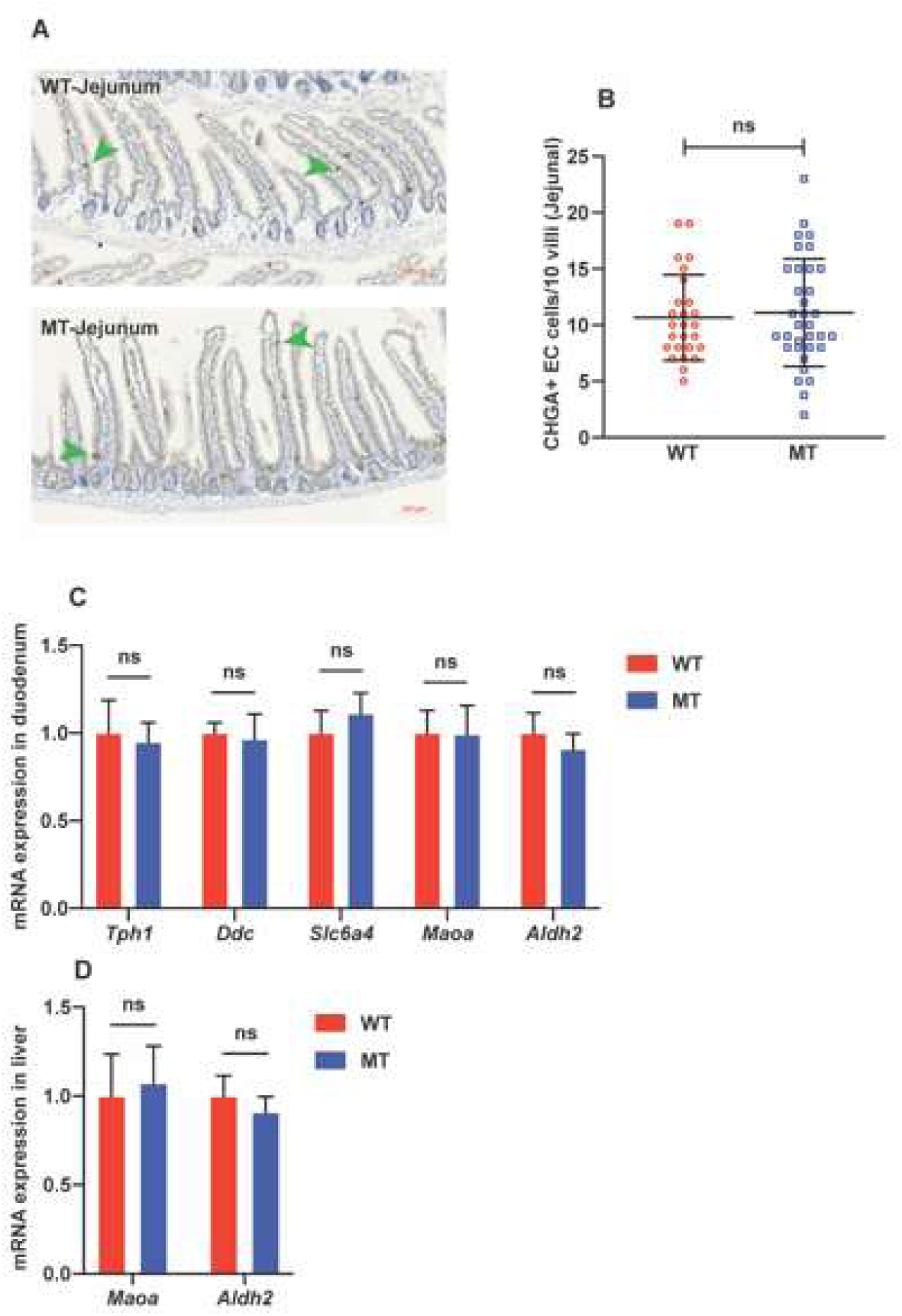
Jejunal serotonin-producing enterochromaffin cell numbers and serotonin metabolism in SI and liver in WT and MT mice. **A** Representative images of jejunal tissue sections stained with anti-chromogranin A (CHGA) antibody for enterochromaffin cells (in brown, indicated by green arrow heads), in jejuna of WT and MT mice, with haematoxylin-counterstained nuclei (in blue). Scale bar = 100 μm. **B** Quantification of CHGA^+^ enterochromaffin cells (number of cells per 10 villi) in jejuna of WT and MT mice (n=6 per group). **C** mRNA expression of enzymes for serotonin synthesis *Tph1* (tryptophan hydroxylase 1) and *Ddc* (5-hydroxytryptophan decarboxylase), serotonin transporter *Slc6a4*, and enzymes for serotonin metabolism *Maoa* and *Aldh2*, in the duodena of WT and MT mice (n=8 per group). **D** mRNA expression of enzymes for serotonin metabolism *Maoa* and *Aldh2* in the livers of WT and MT mice (n=8 per group). mRNA levels were normalized to the control gene *Hprt1*. Data graphs show means ± S.D. *p* values were calculated with the student’s *t* test. ns denotes not significant with *p* ≥ 0.05 between indicated groups.

**Supplemental Figure 9.**
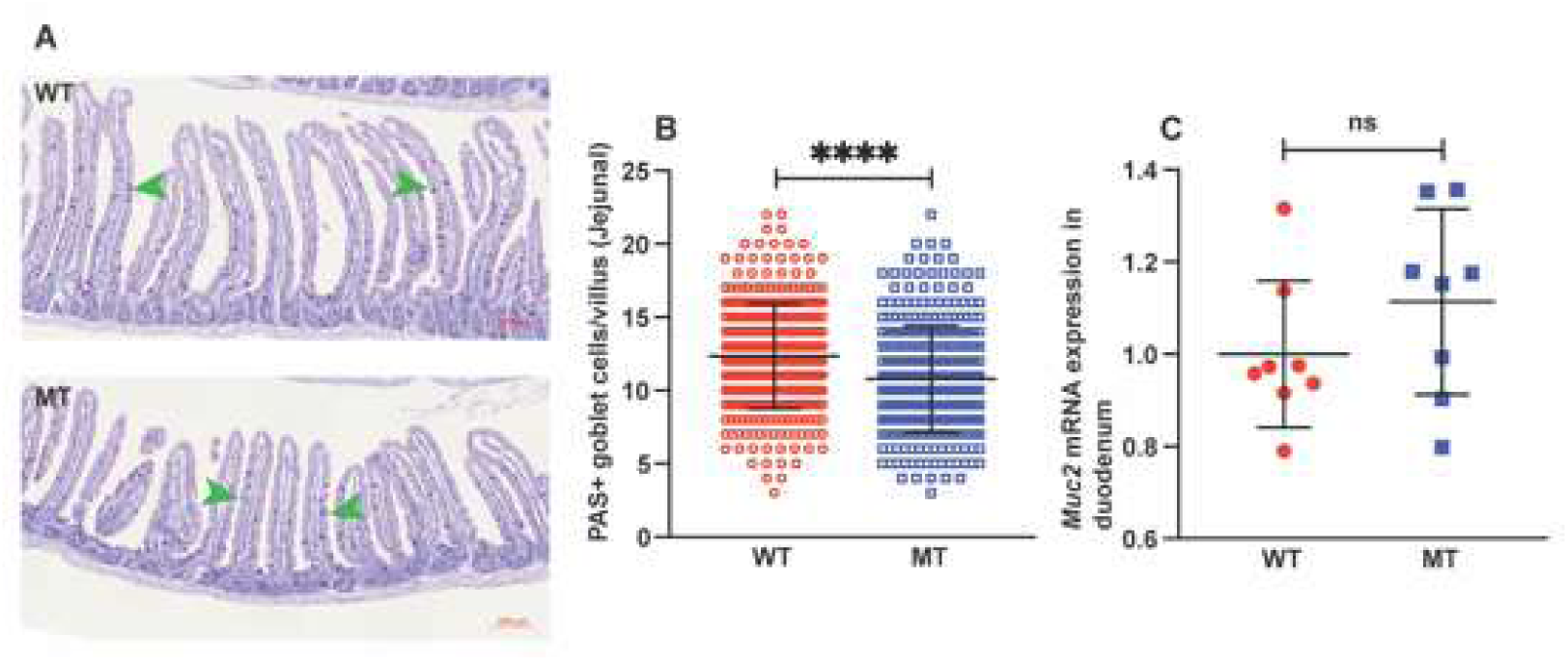
Indole-non-producing *E. coli*-colonized mice contained fewer mucus-producing goblet cells in the small intestines. **A** Representative images of jejunal tissue sections stained with periodic acid-Schiff (PAS) reagent for goblet cells (in deep purple, green arrow heads indicate representative cells), in jejuna of WT and MT mice, with haematoxylin-counterstained nuclei (in blue). Scale bar, 100 μm. **B** Quantification of goblet cells (number of PAS^+^ cells per villus) in jejuna of WT and MT mice (n=6 per group). **C** mRNA expression of mucin 2 (*Muc2*) in the duodena of WT and MT mice (n=8 per group). mRNA levels were normalized to the control gene *Hprt1*. Data graphs show means ± S.D. *p* values were calculated with the student’s *t* test. Statistical significance is presented as \*\*\*\**p* < 0.0001 between indicated groups. ns denotes not significant with *p* ≥ 0.05.

